# A Bayesian Two-Way Latent Structure Model for Genomic Data Integration Reveals Few Pan-Genomic Cluster Subtypes in a Breast Cancer Cohort

**DOI:** 10.1101/387076

**Authors:** David M. Swanson, Tonje Lien, Helga Bergholtz, Therese Sørlie, Arnoldo Frigessi

## Abstract

**Motivation:** Unsupervised clustering is important in disease subtyping, among having other genomic applications. As genomic data has become more multifaceted, how to cluster across data sources for more precise subtyping is an ever more important area of research. Many of the methods proposed so far, including iCluster and Cluster of Cluster Assignments, make an unreasonble assumption of a common clustering across all data sources, and those that do not are fewer and tend to be computationally intensive.

**Results:** We propose a Bayesian parametric model for integrative, unsupervised clustering across data sources. In our two-way latent structure model, samples are clustered in relation to each specific data source, distinguishing it from methods like Cluster of Cluster Assignments and iCluster, but cluster labels have across-dataset meaning, allowing cluster information to be shared between data sources. A common scaling across data sources is not required, and inference is obtained by a Gibbs Sampler, which we improve with a warm start strategy and modified density functions to robustify and speed convergence. Posterior interpretation allows for inference on common clusterings occurring among subsets of data sources. An interesting statistical formulation of the model results in sampling from closed-form posteriors despite incorporation of a complex latent structure. We fit the model with Gaussian and more general densities, which influences the degree of across-dataset cluster label sharing. Uniquely among integrative clustering models, our formulation makes no nestedness assumptions of samples across data sources so that a sample missing data from one genomic source can be clustered according to its existing data sources.

We apply our model to a Norwegian breast cancer cohort of ductal carcinoma in-situ and invasive tumors, comprised of somatic copy-number alteration, methylation and expression datasets. We find enrichment in the Her2 subtype and ductal carcinoma among those observations exhibiting greater cluster correspondence across expression and CNA data. In general, there are few pan-genomic clusterings, suggesting that models assuming a common clustering across genomic data sources might yield misleading results.

**Implementation and Availability:** The model is implemented in an R package called twl (“two-way latent”), available on CRAN. Data for analysis is available within the R package.

**Supplementary Material:** Appendices are available online and include additional Breast Cancer subtyping analysis and model runs, comparison with leading integrative clustering methods, fully general statistical formulation and description of improvements of the Gibbs sampler.

## 1 Introduction

Interest in integrative genomic analysis has grown in recent years as the ability to measure a range of genomic features at reasonable cost has increased [Cava et al., 2015, Huang et al., 2012, Zhang et al., 2012, Speicher and Pfeifer, 2015, Kristensen et al., 2014]. Integrating different genomic data sources is motivated by understanding the data at the level of biological systems, rather than discrete spaces to only be understood in isolation of one another [Ali et al., 2014, Curtis et al., 2012, Kristensen et al., 2014, Myhre et al., 2013, Reiss et al., 2006]. It is apparent that the study of functional genomics benefits from acknowledgement of this interplay between data layers [Sun et al., 2018].

Clustering algorithms have an important role to play in integrative genomics. One example is that disease subtypes often manifest themselves as distinct clusters in one or multiple datasets [Sørlie et al., 2001, 2003, Koboldt et al., 2012]. As awareness of tumor heterogeneity and grows and data granularity improves, one might also begin using these methods to understand the genomic landscape within tumor [Wang et al., 2014, Yap et al., 2012, Park et al., 2010]. Sometimes cluster boundaries are shared across data sources, and sometimes each data source has distinct sets of clusters [Netanely et al., 2016]. Since many of these subtypes are still relatively unknown, unsupervised clustering algorithms have a unique role to play in increasing biological understanding of disease heterogeneity–unsupervised clustering orients itself more towards discovery than those methods which implicitly encode a priori assumptions with class labels.

Integrative clustering approaches have taken different tacks both in terms of underlying assumptions of the biological mechanisms and fitting procedures of the algorithms. An important distinction in the former relates to relative placement of clusters; many models assume that clusters and their boundaries are held in common across data sources and try to best increase statistical power for finding these shared boundaries under this assumption [Shen et al., 2009, 2013, Chalise and Fridley, 2017, Chalise et al., 2014, Chen et al., 2008, Mo et al., 2013]. When common boundaries are assumed, some models additionally assume common scaling across different datasets, necessitating “pan-normalization” of them [Lock et al., 2013, Hellton and Thoresen, 2016]. One disadvantage of the assumption is that even when normalization is done well the discrete nature of the data of certain genomic platforms (eg., methylation data) is in fact not amenable to direct comparison to that which is continuous (eg, RNAseq data). Other models make no assumption about relative placement of cluster boundaries across genomic data sources, but parametrize the model to find common boundaries if they exist [Kirk et al., 2012, Dunson and Herring, 2005, Lock and Dunson, 2013, Gabasova et al., 2017]. These methods additionally tend to draw boundaries in a probabilistic way, such that cluster assignment has a distribution rather than definite label. In some cases, each observation has a single cluster distribution, rather than the observation-dataset pair.

Another distinction in integrative clustering algorithms relates to frequentist, Bayesian, and algorithmic formulations of the models. This distinction has implications on the fitting procedures and their scalability. Frequentist formulations of clustering models have generally been those that assume common cluster boundaries across data sources [Kormaksson et al., 2012, Speicher and Pfeifer, 2015, Shen et al., 2009, Mo et al., 2013]. Algorithms with similar aims that decompose aggregations of data matrices into lower dimension spaces have also been developed [Chalise and Fridley, 2017, Chalise et al., 2014, Chen et al., 2008]. Mo et al. [2017]. A Bayesian model makes the common boundary assumption and additionally encodes sparsity into the model via use of priors on variable inclusion. These models have many attractive properties including power gains for finding cluster boundaries when they exist. An important drawback of these models is that due to the iterative nature of model fitting with algorithms like EM and, in other cases, large matrix inversions often involved, they do not scale well to thousands of covariates from many different genomic platforms. An additional drawback is that the number of clusters must generally be specified a priori based on exploratory analysis or post-fitting checks of model fitness criteria, rendering the fitting process multi-stage (eg, [Mo et al., 2017]). Bayesian models draw probabilistic cluster boundaries and additionally tend to benefit from learning the number of clusters [Kirk et al., 2012, Dunson and Herring, 2005, Lock and Dunson, 2013, Gabasova et al., 2017]. While these approaches often sample closed form conditional posteriors quickly, sometimes iterating involves computationally costly operations (eg, [Lock and Dunson, 2013]).

We propose a Bayesian, unsupervised, integrative clustering model that attempts to combine many of the strengths of approaches outlined above. Our model is based on two sets of cluster assignment variables of each sample in each dataset: the first set follows a priori a multinomial distribution, for each dataset independently, so that all cluster assignment variables share the same prior within each dataset; the second set of cluster assignment variables is such that each such variable has the same (sample dependent) multinomial probability in each datasets. In this case each sample will have a priori the same probability to be assigned to a cluster label in all datasets. The intuition behind this construction is that the first construction allows learning cluster assignments within each dataset, the second follows the sample across all datasets. By conditioning on coherence of these two assignments, we are able a posteriori to learn in a natural way within and across datasets. The final cluster assignments depend on three components: a priori cluster probabilities within-dataset (ie, across observations), a priori cluster probabilities within-observation (ie, across dataset), and the likelihood model (ie, how similar the feature vector is to other feature vectors in the same cluster). With appropriate Dirichlet conjugate priors on cluster probabilities, we obtain conditional posteriors for each observation-dataset cluster assignment variable. We describe a Gibbs sampler implementation which performs inference on all parameters.

Our two-way latent structure model (TWL) bears some resemblance to a fully-parametric, Cluster-of-Cluster Assignments (COCA) approach–cluster parameters are dataset specific (thereby making no unreasonable assumption of common scaling or centering across disparate genomic data platforms), but cluster information is shared across data sources [Koboldt et al., 2012, Hoadley et al., 2014]. Unlike COCA, a cluster posterior for the TWL model exists for each observation-dataset, though depending on how cluster boundaries align across datasets, this cluster posterior may be dataset-invariant. Our method is not a two step approach, where uncertainty of the first step is ignored, but produces a coherent posterior uncertainty quantification.

Our model is more flexible than many integrative clustering models in that we do not assume that the same samples are measured in the different datasets (nestedness of observations across data sources). Our literature review of data integration methods suggests this is a unique feature of our model. The special case of no common samples across data sources simplifies to fitting models on each dataset independently. An additional benefit of the model’s formulation is run-time that scales linearly in the number of features. We also learn the number of clusters. One can influence the amount datasets share information through manipulation of hyperparameters as we demonstrate in simulation.

We propose ways to post-process the estimated posterior probabilities for discovering clustering within datasets and identifying that subset of samples whose cluster assignments may span across all datasets. This last metric allows us to examine enrichment in clinical annotation of those samples with greater pan-dataset cluster assignment correspondence. Though we call this “cluster correspondence” because of the context of our model, its calculation is identical to Kirk et al. [2012]’s “cluster fusing” probabilities.

Our paper is organized as follows: in Section 2 we introduce the model, followed by practical considerations and modifications of it. In Section 3, we describe post-processing metrics and simulation studies to develop intuition. We then perform a data analysis of in-situ ductal carcinoma (DCIS) and invasive (IDC) breast cancer using our TWL model and find 2 copy number clusters, 7 distinct methylation clusters, and confirm the 5 known breast cancer subtype expression clusters [Sørlie et al., 2001]. Posterior metrics reveal little correspondence between these disparate clusterings, suggesting that models assuming common cluster boundaries across our datasets could be missleading. We also find modest stratification of the DCIS and invasive samples depending on data source, indicating that the distinction in tumor state may not be reflected equally in genomic data sources. In the Supplementary Material we include a discussion of our modified Gibbs sampler, comparisons with other, common approaches including iCluster and COCA [Shen et al., 2009], how we address the challenge of label switching, and supporting tables and Figures referenced in-text.

## 2 Methods

### 2.1 Model

To give context to description of the TWL model, we use language assuming a clinical study setting where we seek to cluster subjects on whom we have different genomic data sources. However, the model is equally applicable to a setting in which we seek to cluster genes, on which we have different genomic data sets. We also assume each genomic data source has a common set of subjects and no missing values for notational and conceptual clarity, but formulate the more general model (used for our data analysis), which makes no such assumption, in the Supplementary Material.

Let *Y*_*i,j*_ be a vector of features for observation *i* and data set *j* and of dimension *d*_*j*_. So the *j*^*th*^ data source has information on *d*_*j*_ features. We assume *j* ∈ {1*,…, J*}, for a total of *J* data sources, and *i* ∈ {1*,…, N*}. For example, the first data set (j=1) could be the expression values of *d*_1_ genes, and the second data set (j=2), the copy number variation of *d*_2_ loci.

Consider viewing the *Y*_*i,j*_ vectors of length *d*_*j*_ as the atomic units of our model. In doing so, we can think of the TWL model as having 2 separate axes, rows (each row corresponding to a sample) and columns (datasets). We work with a different clustering of the samples for each data set so that every sample is assigned to *J* clusters. We assume that samples tend to be grouped in the same cluster in the various data sets so that a certain cluster coordination is hypothesized across data sets (see Figure 1).

We start with a model for the clustering within each dataset. The cluster assignment variables corresponding to the *column* (ie, data source) models are

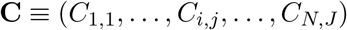

where *C*_*ij*_ ∈ {1, 2,…, *K*} assigns sample *i* in data set *j* to one of *K* clusters, where *K* is the fixed upper bound on the number of clusters. Because the model will tend to find sparse clusterings, *K* is intended to be uninfluential on the number of clusters found. We assume a multinomial prior distribution for the *C*_*ij*_ as

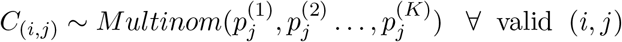

The interpretation of 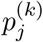 is the probability of a draw of cluster label *k* in dataset *j* for sample *i*.

We emphasize that though *C*_(*i,j*)_ is subscripted by *i* and *j*, the multinomial probabilities are only subscripted by *j*. Therefore, all observations within dataset *j* draw from this dataset-specific *j*^*th*^ multinomial model.

Traditionally, within each dataset, we could model the data vectors *Y*_*ij*_ as a mixture, with one component per cluster and the vector of probabilities 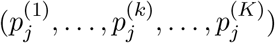 used as mixing parameters. Assuming a parametric model *f* (·) for the density of *Y*_*ij*_, with a cluster dependent parameter 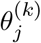, this model would be written as 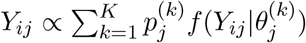.

However, we want to allow and favor alignment in cluster assignments across datasets. For this purpose we propose a new mixture model with different mixing parameters. This is the intuition: in order to help maintain the samples into the same clusters across data sets, we use a second vector of parameters, 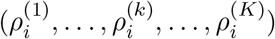, which follow the sample *i* in all data sets. Cluster assignment of a sample in each data set is therefore influenced by both the *p*_*j*_ and the *ρ*_*i*_ parameters. This second parameter *ρ*_*i*_ helps align clusters across datasets because it is the same in each data set. We will combine these two models and use as mixing parameters 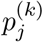 and 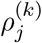 to assign sample *i* to cluster *k* in dataset *j*, as in 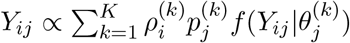.

More formally, we now introduce a second set of cluster assignment variables parametrized by these *ρ* parameters just introduced, which are **R** ≡ (*R*_1,1_,…, *R*_*i,j*_,…, *R*_*N,J*_), where *R*_*ij*_ ∈ {1, 2,…, *K*} assigns sample *i* in data set *j* to one of *K* clusters. We assume a multinomial prior distribution for the cluster assignment variables *R*_*ij*_ corresponding to rows (samples)

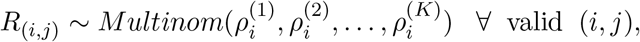

with *K* again the uninfluential upper bound on the number of clusters. Note that though *R* is subscripted by *i* and *j*, the parameters are only subscripted by *i*, the sample id. We can loosely think of the cluster models on the *i*’s, cutting across datasets, as models on the rows of the aggregated data sources if we arranged them next to one another (see Figure 1 as an example). The cluster models for **R** influence cluster alignment along that axis and, in doing so, give across-dataset meaning to cluster labels. This occurs despite cluster density parameters not being directly informed by observations of the same cluster label in different data sets.

It is because cluster labels for the *i*^*th*^ row of dataset *j*, which we denote with *R*_*i,j*_ and *C*_*i,j*_, must agree to produce a coherent and well-defined clustering of samples within each dataset that we condition our model on their equivalence; ie, we condition our likelihood on *R*_*i,j*_ = *C*_*i,j*_ for all valid (*i*, *j*) pairs. Simultaneous use of multinomial models along samples (or “rows”) and within dataset (or “columns”), the resultant necessary conditioning event of *C*=*R* for cluster coherence within each cluster, and the subsequent inferential procedure, are the methodological novelties of the TWL model. This framework results in a joint posterior cluster distribution whose interpretation has granularity at the level of a sample-dataset, rather than just sample. We advocate for this more rich and descriptive, if complex, interpretation in Section 3 below.

**Figure 1:**
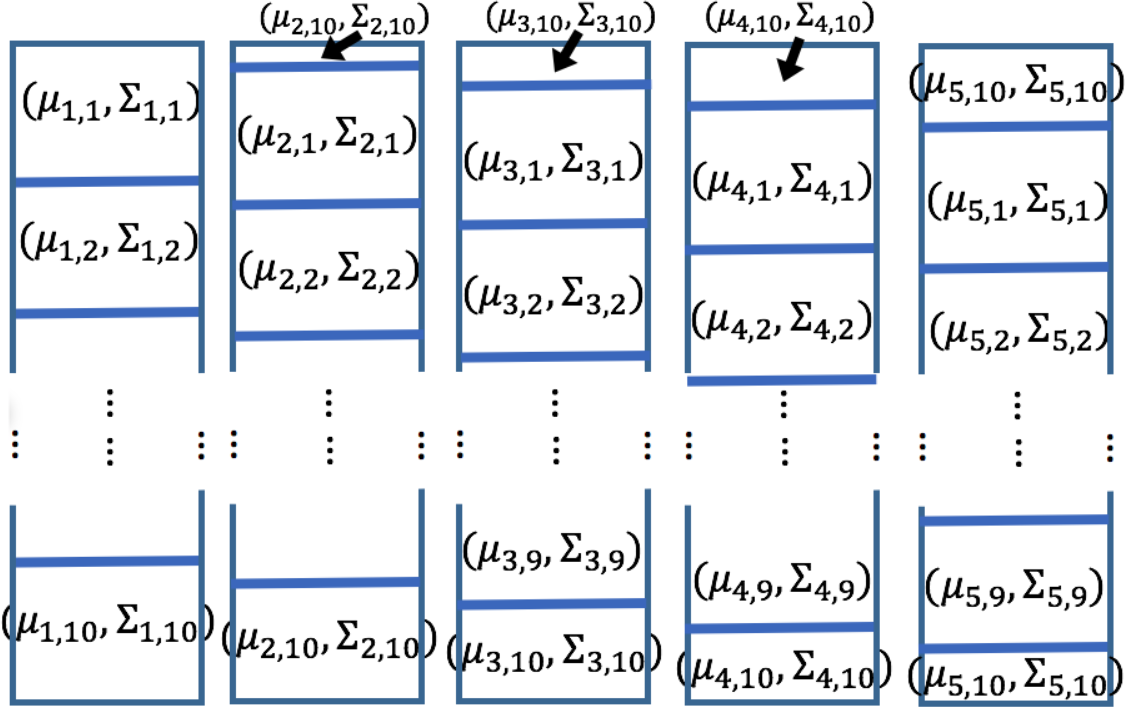
Diagram of progressively misaligned clusters simulation that results in the half-matrix cluster probabilities shown in Table 1. The five long rectangles represent the five generated datasets, each with 200 observations, 10 features, and 10 equally-sized clusters within. Only the first and last few of the 10 clusters can be depicted with boxes in each data set, and the parameters associated with the boxes are those indexing the multivariate normal distributions from which observations in each cluster are sampled. The Σ variance parameters are identical.

As an aside, one can imagine a model similar in spirit whose formulation eschews the need for two sets of latent variables and instead defines its mixing probability as 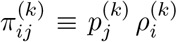, 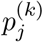 and 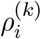 defined as we have them, which in turn parametrize the latent label for sample *i* in dataset *j* and scales density 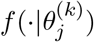. How to efficiently sample such a model is not immediately clear however. Envisioning two sets of latent labels acting along perpendicular axes and conditioning on their equivalence is a way to achieve the clean posteriors we show.

Conditional on *C*=*R*, and on the dataset- and cluster-specific density parameters 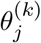 introduced below, *Y*_*ij*_ follows a typical Gaussian mixture model, with cluster probabilities being normalized products of row and column model specific cluster probabilities. For *f* (·) the Gaussian density, the mixture model is

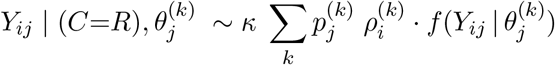

with *κ* the normalizing constant. While we focus on the Gaussian density and thicker-tailed alternatives here [Coretto and Hennig, 2016, 2017, Banfield and Raftery, 1993], one can use any exponential family density with little additional development.

We need to specify the prior model further. The prior probabilities for **p**_**j**_ and *ρ*_*i*_ are

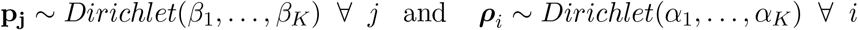

with hyperparameters *α*_*i*_ and *β*_*j*_ constant in *i* and *j* respectively, for which hyperpriors could further be assumed. Instead we choose *α*_*i*_ as a function of the average number of data sets per sample (or simply number of data sets if all samples are present in all data sets and unique within them), and *β*_*j*_ as a function of the average number samples within each data set (or simply the number of unique sample ids if again all samples are present in all data sets and unique within them). As a result, we have *α* ≡ *α*_1_ = *α*_2_ = … = *α*_*K*_ and *β* ≡ *β*_1_ = *β*_2_ = … = *β*_*K*_.

Define the Gaussian cluster density parameters specific to dataset *j* and cluster *k* with 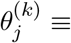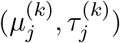, where 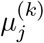 is the mean vector of dimension *d*_*j*_. We assume a priori independence for elements of vector *Y*_*i,j*_ so that precision parameter 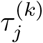 is also of dimension *d*_*j*_. The assumption increases computational efficiency significantly, and filtering correlated, redundant features before analysis makes the assumption reasonable. We generalize beyond the Gaussian likelihood in Section S1.1.2.

We place a Gaussian prior on 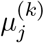 with mean and precision parameters *μ*_0*,j*_ and *ψ*_0*,j*_, respectively. We take an Empirical Bayes approach to choosing these hyperparameters, setting *μ*_0*,j*_ to the mean of *Y*_·,*j*_ (ie, dataset *j*), and the corresponding precision parameter to

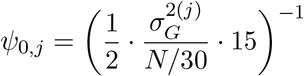

for all *j*, where 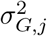 is a vector of variances of the features of *Y*_*·,j*_. Setting *ψ*_0*,j*_ in this way means that if half of marginal variation in data set *j*, 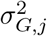, is attributable within-cluster and the number of equal-sized clusters *K* = 30, then there will be 1/15 cluster mean shrinkage to the global mean under such circumstances.

Such a prior also serves a practical purpose: without it, small clusters, especially those with one observation, will be resistant to becoming unoccupied since the mean parameter *μ*_*k,j*_ will tend to be close or identical to the corresponding (few) observation(s) and thus inflate the likelihood. By shrinking *μ*_*k,j*_ to a global mean, especially when the cluster size is small, the posterior of *C, R* | **p**, *ρ*, **C**=**R** will exhibit greater cluster sparsity than that already encouraged by the multinomial priors. In practice, we found simulation and data analysis results insensitive to this choice because distance between peaks in the likelihood makes posterior cluster assignment stable.

With this same goal of convergence, we assumed a common variance parameter *τ*_*j*_(*k*) = *τ*_*j*_ across clusters and one that is considered fixed. Doing so leads to more efficient estimation in the parameter. While the variance across clusters within dataset may not always be constant, since the model has no effective upper bound on the number of clusters, the posterior tends to favor the partition of more heterogeneous clusters into smaller, more homogeneous ones. This characteristic is likely helpful for clincal interpretation.

The full conditional posterior for *C* (which we only consider since we condition on C=R), is

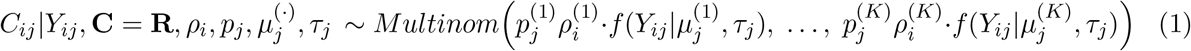

The full conditional posterior for 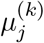 is

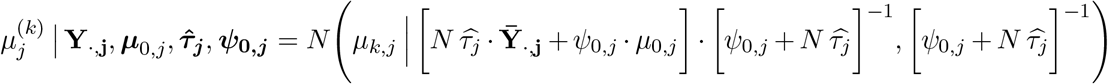

where 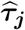 is estimated from the data according to an empirical Bayes approach, assuming 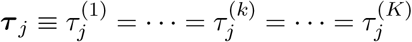.

The full conditional posteriors for *p*_*j*_ and *ρ*_*i*_ are, respectively,

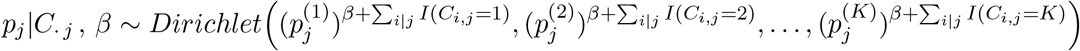

and

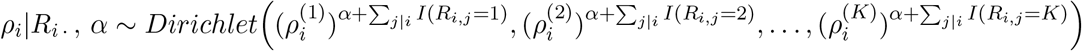

We write the posterior of *ρ*_*i*_ with *R*_*ij*_ to emphasize *ρ*_*i*_’s connection to the “row” multinomial models. However, since *C*_*ij*_ = *R*_*ij*_ for all (*i, j*) by assumption, using *C*_*ij*_ instead is an identical formulation. Additionally, we use the notation *i* | *j* and *j* | *i* to denote valid values of *i* and *j* within strata of *j* and *i*, respectively. We use this notation despite assuming a common set of subject ids for all datasets in this simpler formulation of the TWL model to emphasize that no such assumption is necessary.

We report the general model in the Supplementary Material which assumes dataset-specific sets of sample ids, along with modifications of our Gibbs sampler and generalization of the density function beyond the Gaussian case.

#### 2.1.1 Hyperparameter tuning

Choice of *β* and *α* can be a function of the total number of samples *N* and number of datasets *D*, respectively, and the specified maximum number of clusters. If we consider *β*, increasing its value “dilutes” the effect of the realized cluster distribution on the posterior of **p**_**j**_, and subsequently *C*_*·j*_, by increasing the unnormalized probabilities of all labels equally. Doing so leads to less sparse clusterings within dataset as the likelihood term in the posterior for *C*_*·j*_ is weighted relatively more and therefore pushes heterogeneous observations into their own cluster rather than sharing one. One can calculate the degree by which we dilute the effect of realized latent labels with *β* by considering the sample size *N* (ie, dimension of *C_·j_*) and upper bound on the number of clusters, *K*: since *N* observations have latent labels *C*_*ij*_, adding *β* to each of *K* categories of **p**_**j**_ makes those *N* labels have *N/*(*N* + *K* ∗ *β*) of their original influence on the within dataset clustering.

An analogous calculation and similar reasoning holds for *α* and its effect on *ρ*_*i*_ and *R_i·_*, though in this case one uses *D* (ie, dimension of *R_i·_*), the number of datasets in the calculation. Because *D* will tend to be much smaller than *N*, *α* will likewise tend to be chosen smaller than *β*. The special case of *α* → ∞ leads to no cluster information sharing across datasets because realized latent labels are diluted to no effect on the posterior, effectively fitting independent mixture models on each dataset. Since *K* is intended to be chosen as uninfluential on the number of clusters found, by selecting *α* and *β* so that the proportions *N/*(*N* + *K* ∗ *β*) and *D/*(*D* +*K* ∗*α*) are constant for changes in *K*, one can keep these hyperparameters’ influence on model posteriors constant. We discuss choice of *α* and *β* more in the Supplementary Material.

## 3 Results

### 3.1 Simulation

We generated several different data sets to demonstrate different properties of the TWL model. First, we generated 2 different data sets with nested cluster patterns across data sets (See Figures 2 and 3). Second, to demonstrate how common cluster sharing across data sets can be used to learn structure of cluster patterns in the data sets, we generated “progressively misaligned” clusters, whose diagram can be seen in Figure 1. These simulations illustrate how cluster assignment in one data set influences that in other ones, depending on alignment pattern. They also resemble real genomic data sets whose cluster patterns are not often perfectly aligned across genomic datasets and which exhibit nestedness patterns to some degree [Netanely et al., 2016]. For example, for RNAseq and CNA tumor data on a common set of subjects, clusters defined solely by RNAseq and solely by CNAs would likely not align. Additionally, within, say, a CNA cluster, there very well may be multiple RNAseq clusters. Ability to find such structure is an emphasis of our model.

**Figure 2:**
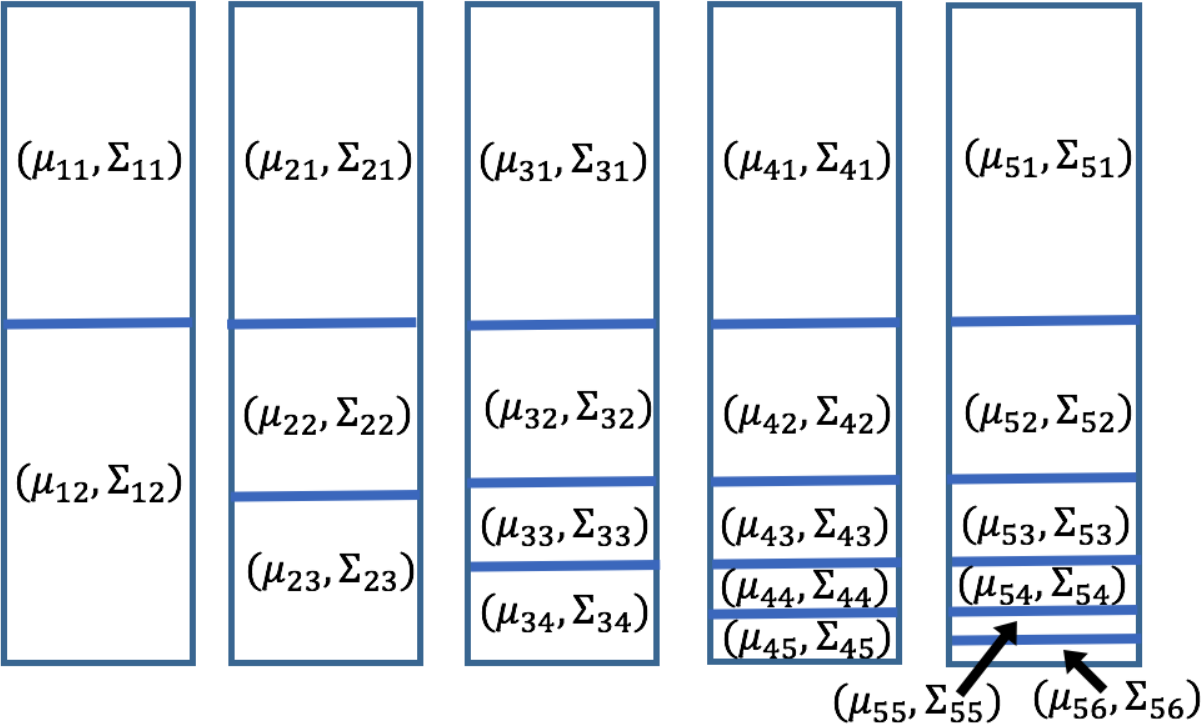
Diagram of nested clusters simulation. The five long rectangles represent datasets, each with 200 observations and 10 features, and boxes within them the unique clusters composing the datasets. The parameters in each box index the multivariate normal distributions used to generate samples in each cluster. As one moves from left to right along datasets, one new cluster is added relative to the previous dataset by halving the bottommost cluster in that dataset. The Σ variance parameters are equivalent.

**Figure 3:**
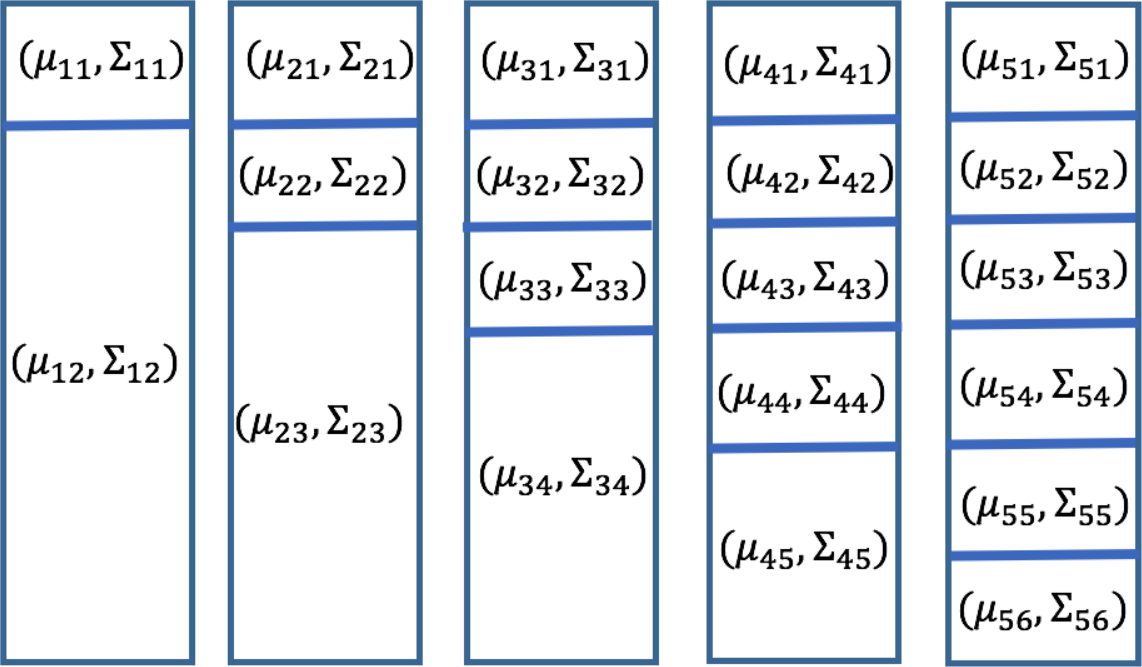
Diagram of a clusterings in the nested clusters simulation. The five long rectangles represent datasets, each with 200 observations and 10 features, and boxes within them the unique clusters composing the datasets. The parameters in each box index the multivariate normal distributions used to generate samples in each cluster. As one moves from left to right along datasets, one new cluster is added relative to the previous dataset. The Σ variance parameters are equivalent.

#### 3.1.1 First nested scenario

We see in Figure 2 our first nested data scenario, a collection of five data sets. Each data sets consists of 200 observations, each with 10 features. In each of the five data sets, observations in blocks denoted in the figure are drawn from the same multivariate normal distribution. The multivariate normal distributions corresponding to the different blocks have the same covariance matrix but different means. Means were generated randomly with each sampled element in the mean vector having a standard deviation of 1.4. The standard deviation of observations from respective means was 1 in all 10 feature dimensions, with features generated independently. We make this independence assumption in simulation and later analysis to significantly increase computational efficiency. The assumption is valid in analysis if data is processed so that highly correlated features are excluded. Moving from left to right over the 5 data sets in Figure 2, the number of clusters varies from 2 to 6, with each additional cluster in the next data set resulting from splitting the bottom cluster in the previous data set in half and introducing slight misalignment. Such a telescoping pattern helps us understand how well our model identifies the small clusters in the right-most data sets and how common cluster labels are shared across data sets. There is a trade-off between fitting larger, “umbrella” clusters, with a single cluster label, and using multiple, different labels for the “enclosed” clusters in successive data sets because of information sharing across them all. Analyzing how our model resolves these trade-offs yields information on relative signal clarity and size of clusters, and their alignments.

#### 3.1.2 Second nested scenario

Figure 3 shows our second nested data scenario with five data sets, each 200 observations by 10 features. Now all clusters added in subsequent data sets are of the same size. Such a pattern allows us to again examine the effect of nested clusters in posterior cluster labels, but no one cluster becomes especially small in the furthest right data sets, unlike the first nested scenario.

#### 3.1.3 Progressively misaligned scenario

Figure 1 shows a set of five data sets of the same dimension as those above exhibiting no nestedness in its cluster patterns. In this scenario, however, clusters are progressively misaligned. The first data set consists of 10 clusters each consisting of 20 observations. The next data set to the right also has 10 clusters, though those clusters are misaligned by 2 observations (10% of the size of each cluster). The next data set to the right again has 10 clusters, again misaligned by 2 observations as compared to the second data set. This pattern continues until the 5th dataset is almost entirely misaligned as compared to the first data sets, by 8 observations total. Data sets were generated in this way in order to examine how posterior cluster labels are held in common in less (e.g., the third and fourth data sets) and more (the first and fifth data sets) cluster-misaligned datasets.

### 3.2 Posterior interpretation

Unlike many other integrative clustering models, the TWL model yields cluster assignments for each sample in each dataset. One can therefore examine for each observation a measure of cluster membership across datasets, which we call “cluster correspondence”. We propose this measure and four related metrics to understand clustering patterns in our data sets.

#### Metric 1 (“Within”)

This metric looks at cluster co-assignment of observations, within datasets. We calculated the matrix of elements

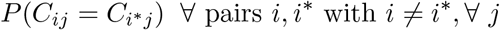

which we estimate by the proportion of samples in our chain after burn-in where observations *i* and *i*\* have a common cluster label. These proportions result in *J* symmetric *N* × *N* matrices. Figures 4 and 5 show two examples of heatmaps of such output matrices.

**Figure 4:**
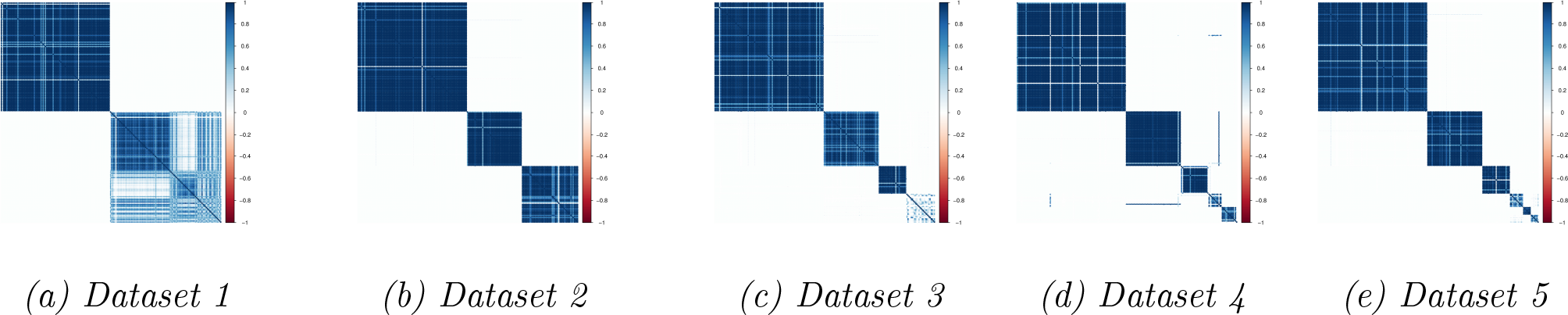
Metric 1 for the 5 datasets from the first nested cluster simulation, whose diagram is Figure 2. The heatmap depiction of the metric makes clear that simulated clusterings are generally identified with high fidelity. The last clusters in (a) and (c) are not identified as clearly, likely related to fracturing of clusters in adjacent datasets.

**Figure 5:**
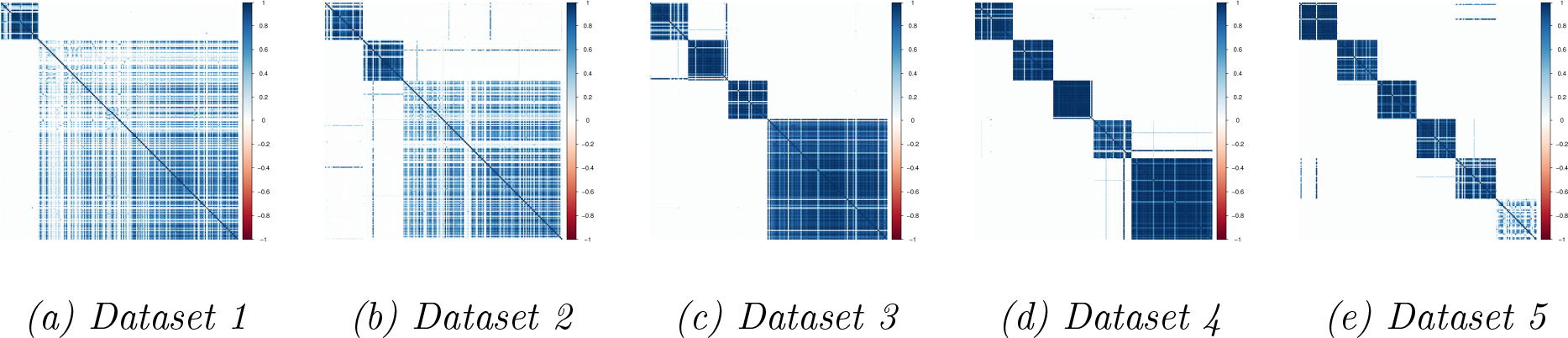
Metric 1 for the 5 datasets from the second nested cluster simulation, whose diagram is Figure 3. The heatmap depiction of the diagnostic shows that there is some difficulty identifying the largest clusters in (a) and (b), and the bottommost cluster in (e). At least in the case of the large clusters, this is likely due to cluster division in adjacent datasets per the simulation.

#### Metric 2 (“Between”)

This metric looks at cluster assignment between datasets, within observations. We calculate the matrix of

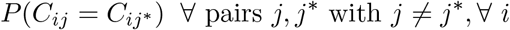

There are *N* such matrices, one for each observation, each of dimension *J* × *J* with *J* ∗ (*J* − 1) unique elements. If we consider the more general TWL model where samples are not assumed nested across datasets, some elements in a subset of these matrices may be NA if the corresponding observation does not have data on both data sources associated with that element. The interpretation of an element of a particular matrix is the proportion of common cluster assignments for that pair of data sets for that observation.

#### Hierarchical clustering

We perform hierarchical clustering on each of the *N* × *N* matrices from Metric 1. The recommended workflow is to first examine heatmaps of the matrices generated in Metric 1, determine the number of clusters, and then cut the dendrogram at the appropriate place to generate that number of clusters and make a determination of cluster membership if that is needed and desired. We make additional suggestions about this post-processing analysis in Section 3.

#### Between-dataset cluster correspondence

Consider the *N* matrices from Metric 2: we can average element-wise over all such *N* matrices. The interpretation of entry (*j, j\**) in the resulting symmetric *J × J* matrix with elements in [0, 1] is the degree to which cluster boundaries are similar in datasets *j* and *j\**. Higher numbers indicate more similar boundaries.

The best example of the use of this measure is for the misaligned cluster simulation (see Figure 1) and demonstrates that the more clusters do not align, the lower are elements in the matrix. Indeed, we observe a decreasing cascade of proportion of common clusters for more “distal” datasets, as the misalignments in datasets increases.

#### Sample-specific cluster correspondence

Consider again the N matrices from Metric 2: we can average over the *J* (*J* − 1) unique elements in each matrix, resulting in a length *N* vector. The interpretation of the entry corresponding to observation *i* in the vector, between 0 and 1, is the degree to which cluster membership is similar across the *J* datasets for that observation, what we call sample-specific “cluster correspondence”. A higher number indicates greater common cluster membership across datasets for the observation. One can also calculate cluster correspondence just for subsets of datasets for more specific hypotheses.

### 3.3 Simulation results

We show the PSMs for our 2 nestedness simulations in Figures 4 and 5. The figures reveal that the TWL model can identify the clusters with a high degree of fidelity. As one might expect from the telescoping nested scenario, the cluster labels in the lowermost block from the first dataset in that simulation scenario are not as stable as the uppermost block, due to the “splintering” of clusters in subsequent datasets and across-dataset influence of cluster membership. One can see a similar phenomenon in the smaller cluster blocks of (c), (d), and (e), and also the larger blocks of Figure 5 (a) and (b) – while their outline and therefore recognition is clear, common labelling of those observations is not as stable, as should be.

We developed the misaligned cluster simulation in part to examine between-dataset cluster correspondence, which is shown in Table 1 for two different values of the *α* hyperparameter. In both (a) and (b), we see decreasing values in across-dataset cluster correspondence as one moves away from the diagonal, in a Toeplitz or auto-regressive pattern as one would expect. We also see the influence of *α* on this measure and again recommend some normalization of that parameter with respect to the total number of dataset used in analysis.

**Table 1:** Tables of the between-dataset cluster correspondence for the misaligned cluster simulation, shown in Figure 1, for different values of α. In either table, entry j, j* is the average cluster overlap of datasets j and j*. One expects more cluster overlap when clusters align to a greater degree, as they do in adjacent datasets according to Figure 1. For either value of α, as one moves further from the diagonal, one notices a cascading decrease in common clusters across data sets as expected. One also observes larger numbers in b) as compared to a), due to the influence of higher values of α diluting the cluster label information sharing across datasets.

|  | dat1 | dat2 | dat3 | dat4 | dat5 |
| --- | --- | --- | --- | --- | --- |
| dat1 | 1.000 | 0.392 | 0.364 | 0.230 | 0.201 |
| dat2 | 0.392 | 1.000 | 0.461 | 0.272 | 0.230 |
| dat3 | 0.364 | 0.461 | 1.000 | 0.314 | 0.299 |
| dat4 | 0.230 | 0.272 | 0.314 | 1.000 | 0.247 |
| dat5 | 0.201 | 0.230 | 0.299 | 0.247 | 1.000 |

b) Using $\alpha = 0.4$ for the misaligned cluster simulation.
|  | dat1 | dat2 | dat3 | dat4 | dat5 |
| --- | --- | --- | --- | --- | --- |
| dat1 | 1.000 | 0.744 | 0.690 | 0.643 | 0.565 |
| dat2 | 0.744 | 1.000 | 0.773 | 0.717 | 0.640 |
| dat3 | 0.690 | 0.773 | 1.000 | 0.838 | 0.745 |
| dat4 | 0.643 | 0.717 | 0.838 | 1.000 | 0.869 |
| dat5 | 0.565 | 0.640 | 0.745 | 0.869 | 1.000 |

We developed the nestedness cluster simulation in part to examine sample-specific cluster correspondence, shown in Figure 6. The horizontal bar chart in (a) is the observation specific cluster correspondence. We observe greater cluster correspondence in observations belonging to clusters in the first dataset that are not split in subsequent data sets. The lowermost observations, whose cluster membership in the left-most datasets are split frequently into smaller clusters in subsequent right-most datasets, have much smaller cluster correspondence measures.

**Figure 6:**
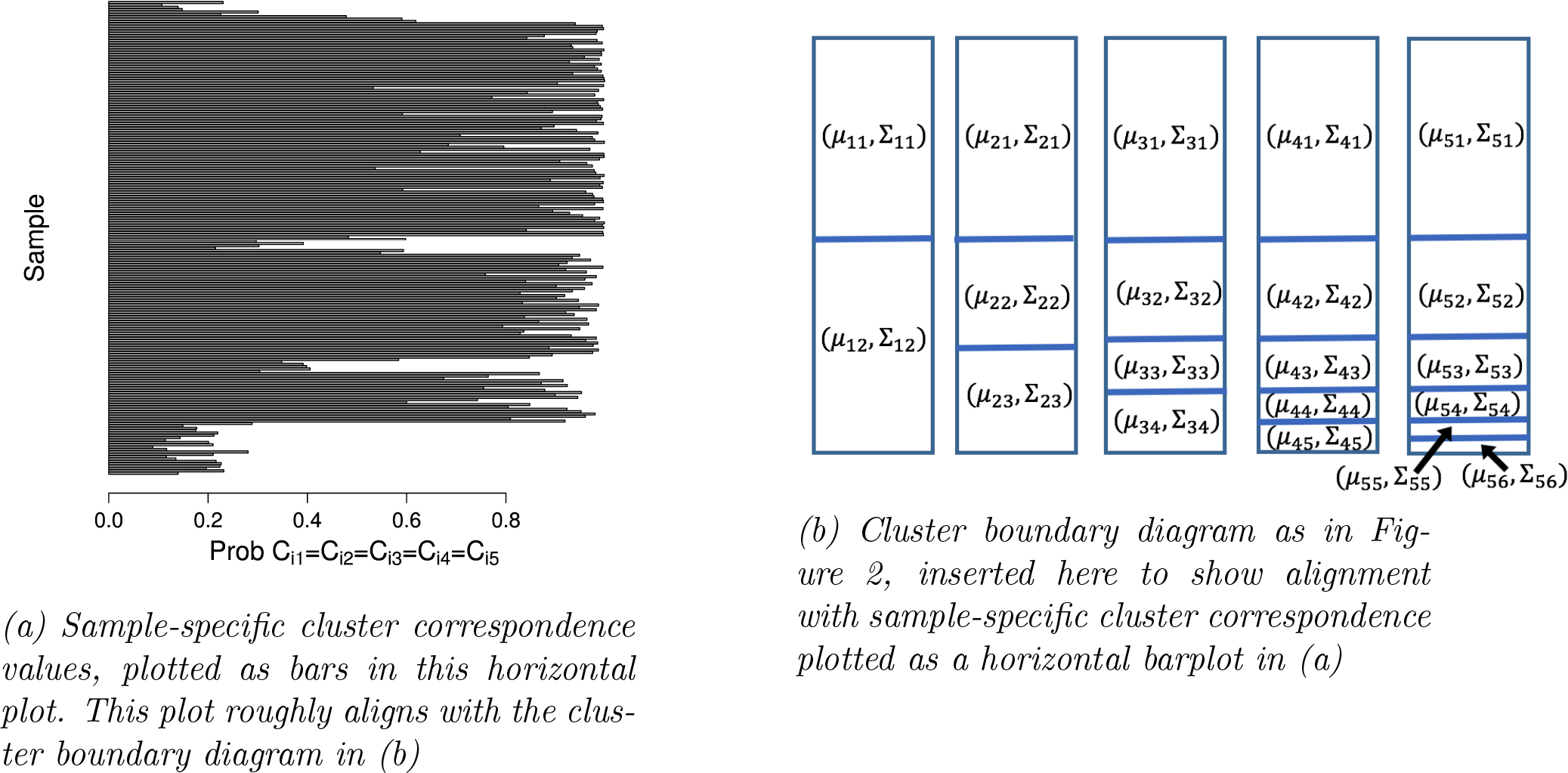
Sample-specific cluster correspondence for the telescoping nestedness simulation

We also calculated the sample-specific cluster correspondence on the misaligned cluster scenario, shown in Figure 7. We observe greater cluster correspondence in those observations whose own row is less likely to be crossed by a cluster boundary in one of the datasets. Since this feature varies by observation and all clusters in all datasets are the same size, one observes growing and shrinking peaks of the cluster correspondence measure as one moves down the horizontal barplot (Figure 7 (a)).

**Figure 7:**
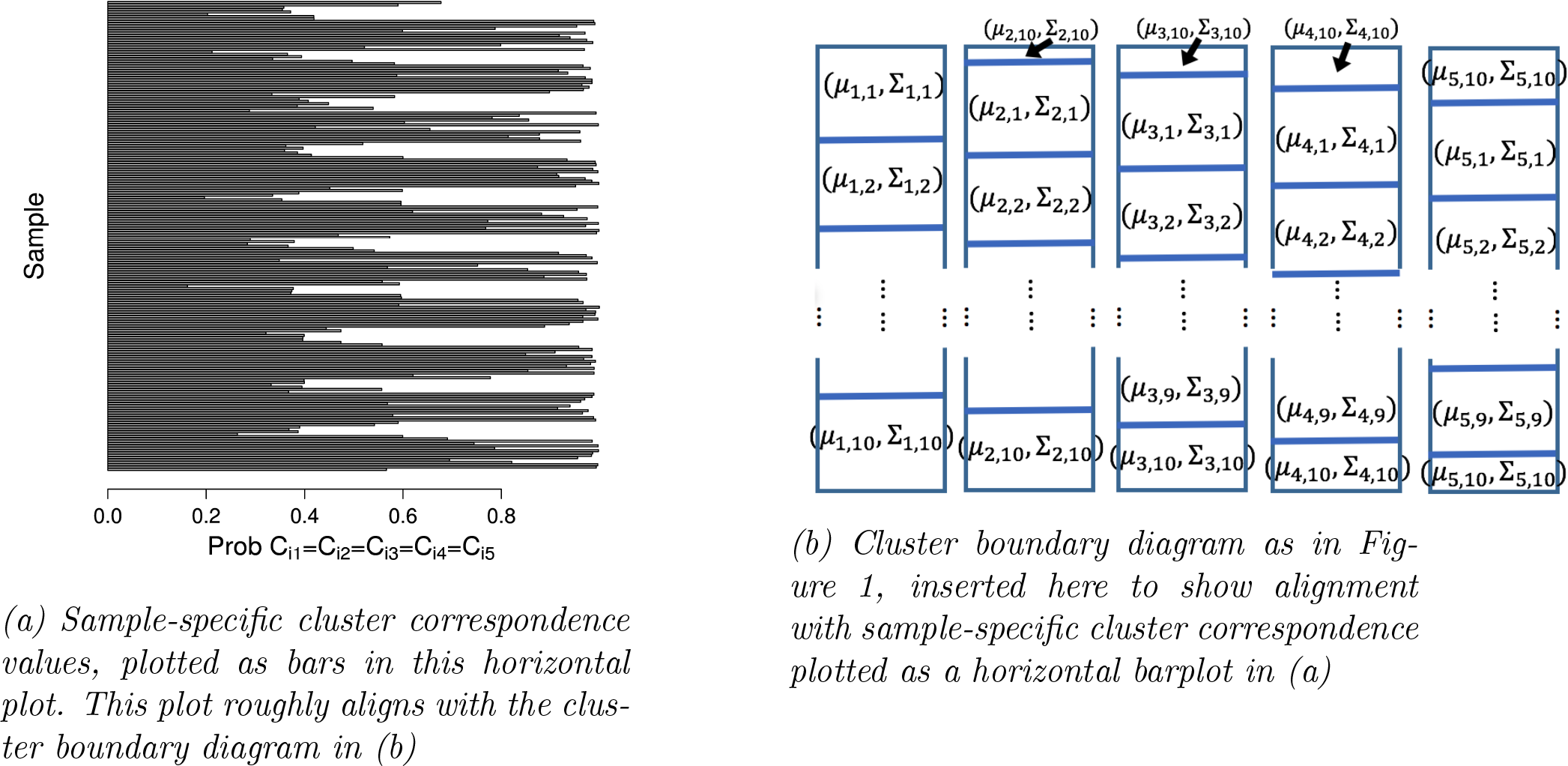
Sample-specific cluster correspondence for the misaligned cluster simulation

### 3.4 Breast cancer subtyping in DCIS and IDC tumors

#### 3.4.1 Preprocessing

We performed an analysis of ductal carcinoma in-situ and invasive breast cancer tumors comprising all intrinsic subtypes coming from a cohort of Norwegian and Italian women. The cohort includes 370 women, 57 in the DCIS tumor state, and 313 IDC, and the data has been described in greater detail elsewhere [Lesurf et al., 2017, Muggerud et al., 2010]. We used Agilent 60K microarrays to measure gene expression on 370 observations and 21887 genes from this cohort. Quantile normalization was performed, and probes collapsed over genes by mean value [Amaratunga and Cabrera, 2001].

We used the Illumina 450K array to measure methylation of CpG sites on 314 of the 370 women, normalized according to the method described in Touleimat and Tost [2012]. We set as missing probes whose detection values had p-values above 0.05. Probes targeting a given CpG were all removed if more than 25% of probes were denoted as missing and performed imputation on those remaining with k-nearest neighbors. Probes were grouped by gene with flanking regions of 50kb, and we collapsed the data to gene-level granularity by using the score according to the first principal component [Wilhelm-Benartzi et al., 2013]. Collapsing the data in this way led to better separation of the DCIS and Invasive tumor states in penalized regression models and so was likewise used to better distinguish these states in our clustering model.

We had copy number alteration data on 338 observations and 19089 genes, the women being a subset of the 370 on whom we had expression data and not a superset of those on whom we had methylation data. We used the SNP 6.0 array and GC-content corrected raw log ratio values. Segmentation was performed using the PCF algorithm in the R package *copynumber* [Nilsen et al., 2012]. We collapsed to gene-level data by determining which copy number segment overlapped the gene most [Curtis et al., 2012]. The Supplementary Material includes model runs using uncollapsed probes from promoter regions (see Figure S2 for example).

#### 3.4.2 Analysis

We performed TWL analysis in the statistical programming language R [R Core Team, 2017]. We ran our Gibbs Sampler for 4000 iterations for different feature draws from the approximately 20000 genes available, in order to investigate reproducibility. Our datasets consisted of approximately 1790 of the same genes from all genomic data sources. We used *α* = 0.4 and *β* = 7 as hyperparameters and used *K* = 30 as our cluster upper bound. Using these hyperparameters means that realized latent label information sharing is diluted ≈ 335/(35 + 7 * 30) ≈ 60% and ≈ 3/(3 + 0.4 * 30) ≈ 20%, 335 chosen as an average of the number of observations over our 3 datasets. We ran also parallel, independent chains on identical draws to assess convergence and Monte Carlo error. Convergence of cluster membership occurred relatively early, though we considered the first 2000 iterations burn-in. During the burn-in phase, we used a warm start strategy for the MCMC to speed convergence of datasets with many outliers. We additionally lower-bounded Gaussian tail densities to robustify the mixture model to clusters of outliers and improve mixing [Coretto and Hennig, 2016, 2017, Banfield and Raftery, 1993] and describe both of these modifications in the Supplementary Material.

PSMs and corresponding heatmaps were generated to give Figure 8. We performed hierarchical clustering analysis on these heatmaps, manually inspected the tree, and cut it at a height so that approximately 95% of the samples had been grouped with at least one other observation (Figure S9). Of those which had been grouped, we gave an identifying label to the M largest clusters (where *M* was 5, 2 or 3, and 7, based on inspection of the expression, CNA, and methylation heatmaps, respectively), and the rest grouped into a more heterogeneous cluster we called “unknown”. The idea behind such a procedure was to label those observations clearly belonging together in the same cluster as such, and to keep those more heterogeneous observations in their own separate group to not dilute signal clarity. The interpretation of placement into the cluster unknown category is not “this sample belongs in a cluster, but we don’t know which one.” Rather, it means “this sample is very different from all the other samples, including those with the clus unknown label, and so we cannot give it a cluster label.” The heterogeneity of samples with this label is especially evident with the Basal subtypes’ CNA data and high genomic instability index (see Figure S3 in the Supplementary Material).

One can use different thresholds for *M* and the tree height depending on proportion of observations falling into a clear cluster. For example, the PSM for CNA suggests that there is a greater proportion of observations with unknown cluster membership, and so we used a dendrogram height translating to approximately 90% of samples already grouped with at least one other observation.

**Figure 8:**
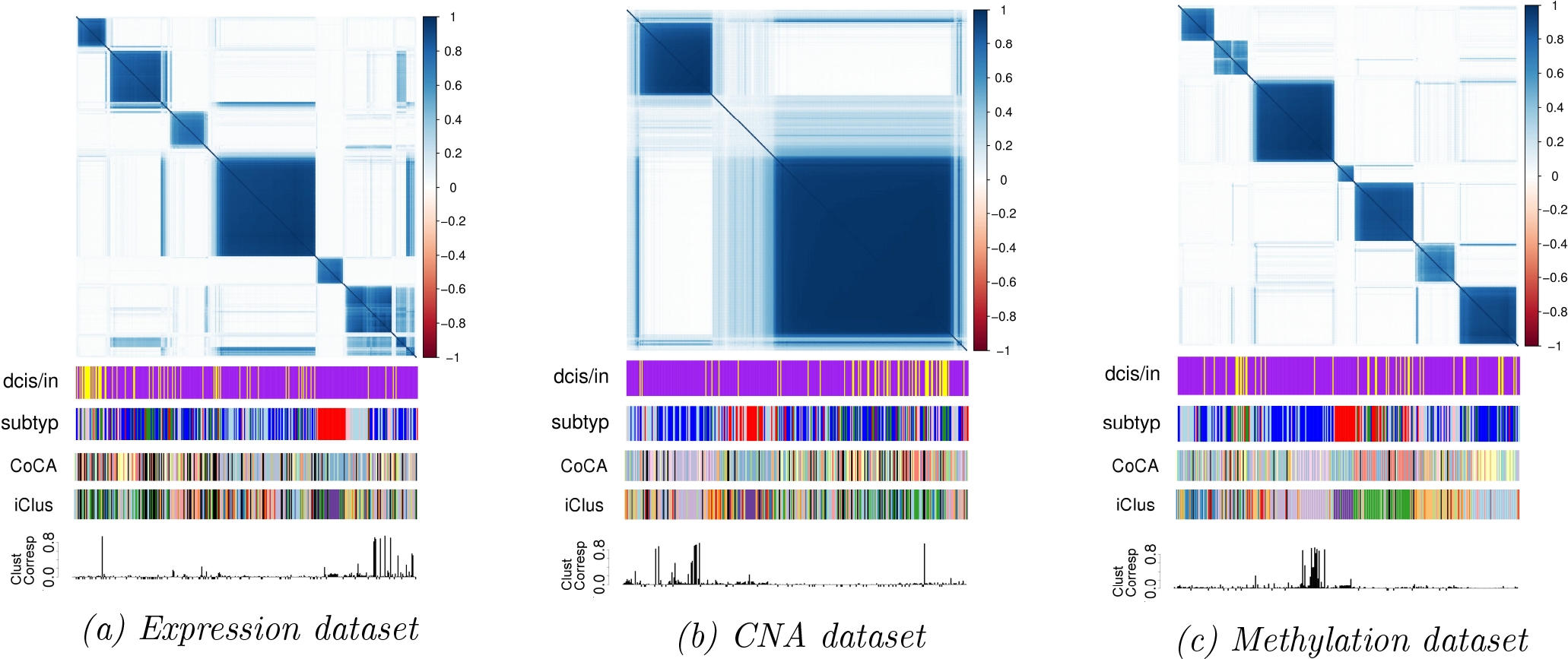
Metric 1 for our breast cancer analysis. There is a high degree of alignment between the known breast cancer subtypes and the clustering for Expression (a). CNAs (b) and Methylation (c) reveal 2 and 7 distinct clusters, respectively. Encoded in our model is cross-dataset cluster assignment influence, though in this analysis interestingly there is relatively little correspondence across datasets. The figures are annotated intrinsic subtype (blue: Lum A, sky blue: Lum B, green: normal-like, pink: Her2, red: Basal), DCIS/invasive tumor state (purple: Inv, yellow: DCIS), and sample-specific cluster correspondence (negative values indicate missing values in the sample in some sources). The Basal intrinsic subtype is the most distinct and compact in each datatype clustering. We also include results from the iCluster and COCA analyses, described in the Supplementary Material.

We calculated the between-dataset cluster correspondence on two different feature draws from our datasets (Table 2), indicative of cluster boundary alignment between data sources as a whole, and found little suggestion of significant boundary overlap in both cases. This stands in contrast to scenarios like our simulations (e.g., Figure 2). Despite the different draws from the data, there was stability in estimation of the measure.

**Table 2:** The between-dataset cluster correspondence with 95% credible intervals. The interpretation of an element in the table is the average cluster overlap between the corresponding dataset pair. There is relatively similar cluster overlap between all pairwise comparisons of genomic datatypes.

|  | Expr | CN | Methyl |
| --- | --- | --- | --- |
| Expr | 1.000 | 0.158 (0.140,0.175) | 0.193 (0.173,0.213) |
| CN | 0.158 (0.140,0.175) | 1.000 | 0.218 (0.202,0.234) |
| Methyl | 0.193 (0.173,0.213) | 0.218 (0.202,0.234) | 1.000 |

We found considerable stability in clustering between parallel MCMC chains (Tables S2 and S3), and relatively high stability in cluster assignment (Tables 3 and S1). In the case of either draw from the data, cross tabulations of at least expression-based cluster membership and breast cancer intrinsic subtypes (PAM50) showed that misclassification, where it occurred, mainly was between more closely related subtypes (e.g., Luminal A and Luminal B, see Table S4, [Sørlie et al., 2003, Parker et al., 2009]). While there is some correlation between CNA or methylation clusters and subtypes, it is not as strong (Table S5). For expression and methylation datatypes, the most distinct subtype was Basal, with perfect specificity and good sensitivity for clusters discovered by our model. It was also most distinct in the CNA data, but in a different way – these samples could not be clustered due to their heterogeneity and therefore are disproportionately represented among samples with the “unknown” cluster label (Table S12). We discuss this result in the Supplementary Material and relate it to Basal samples’ elevated genomic instability index (see Figure S3). Apart from this, samples with the cluster unknown label represented subtypes relatively evenly and were a small proportion of the total samples.

**Table 3:** Post-processed cluster labels for different feature draws from the expression dataset.

|  | clust 1 | clust 2 | clust 3 | clust 4 | clust 5 | clust unknown | Sum |
| --- | --- | --- | --- | --- | --- | --- | --- |
| clust 1 | 2 | 52 | 0 | 6 | 10 | 15 | 85 |
| clust 2 | 0 | 0 | 0 | 0 | 0 | 25 | 25 |
| clust 3 | 4 | 0 | 0 | 102 | 7 | 4 | 117 |
| clust 4 | 0 | 1 | 36 | 1 | 6 | 0 | 44 |
| clust 5 | 24 | 9 | 3 | 1 | 15 | 2 | 54 |
| clust unknown | 3 | 2 | 2 | 4 | 13 | 21 | 45 |
| Sum | 33 | 64 | 41 | 114 | 51 | 67 | 370 |

We discuss the results of the iCluster and COCA analyses in the Supplementary Material, but have included annotation from these analyses in Figure 8. We say briefly that because both methods assume common cluster boundaries across all data sources, the degree to which consensus clusters reflect those of the particular data source varies considerably.

In general cross tabulations of DCIS and Invasive tumor state with posterior cluster labels from the different data sources was not as compelling as that for expression and intrinsic subtypes. An important exception however was a relatively small expression cluster that contained many of the DCIS samples (Table S13). We additionally found that nearly all DCIS tumors were contained by the large CNA cluster identified in our analysis (Table S9). The second, smaller cluster identified in the CNA analysis contained almost entirely invasive tumors. There was a similar finding in the methylation dataset where DCIS tumors were significantly overrepresented in one of the clusters, though a small one in this case.

Because our between-dataset cluster correspondence measure suggested relatively little cluster overlap between datasets on the whole, sample-specific cluster correspondence becomes more interesting: subsets of samples with greater correspondence can be enriched for clinical annotation. We calculated sample-specific cluster correspondence on expression and CNAs and examined the 50 observations with the largest such cluster correspondence. We observed enrichment in the Her2 subtype and DCIS tumor state as compared to the marginal distribution of these variables. These features were observed across parallel MCMC chains and different draws from the data and so appear as robust findings (see Tables S7 and S10). We found similar enrichment in those posterior cluster labels that roughly correspond to the Her2 subtype and DCIS, which may be considered proxies for these types. In either case, there is an important caveat in that both Her2 and its proxy cluster label, and DCIS and its proxy cluster label, are almost entirely contained in the very large CNA cluster one sees in Figure 8b. Thus, this enrichment may be an inevitable result of that cluster’s size and a large proportion of observations with those annotations falling within it. However, the normal subtype is also almost entirely found in this large CNA cluster, but we observe no such enrichment in it when looking at expression-CNA cluster correspondence (Table S12). The finding should therefore be considered with caution, though not discounted.

We observe a lack of cluster correspondence between CNA and methylation clusters among the Basal subtype in particular, across parallel MCMC chains and draws from the data (Table S11). The interpretation of this phenomenon is difficult, in part because of the Basal subtype’s aforementioned presence in the large CNA cluster. One hypothesis is that the alignment of the Basal observations between the expression-methylation and expression-CNA pairs is higher than under the null hypothesis of independent clusterings on each dataset, but on disjoint sets of Basal observations, pushing the alignment for CNA-methylation low, as observed. However examination of the data suggests that this hypothesis likely does not fully explain the lack of CNA-methylation correspondence observed in the Basal subtype. Another unexpected feature of the Basal subtype was its presence in primarily only 2 methylation clusters (Figure S9c).

When looking at pan-genomic cluster correspondence (that measure shown for our simulations in Figures 6a and 7a), we observed no enrichment in subtypes or tumor state. Since we observe enrichment for specific pairs of data sources, the lack of cluster correspondence on all three is either because the enrichment we do see does not carry over to the excluded source, or that there is some bias in types of samples on whom certain data platforms could be measured.

## 4 Discussion

We developed an innovative integrative clustering model called TWL in which we envisioned the existence of multinomial models on clusters along both the “column” (or dataset) and “row” (or observation) of the aggregated datasets. For model coherence, we conditioned on the event that these two otherwise disparate sets of clusterings were equal to one another, resulting in closed form posteriors. The model shares cluster assignment information across datasets, giving across-dataset meaning to the labels, though fits cluster parameters within-dataset. The model learns the total number of clusters with the analyst specifying an upper bound, and posteriors exist on the level of the sample-dataset pair, rather than only on sample which leads to a loss of information. Run time of cluster estimation scales linearly in the total number of covariates in the datasets being integrated.

We argue that forcing a clustering on samples, invariant to data source, seriously oversimplifies diseases subtyping. In our analysis each observation appears best characterized by its full set of clusters, one for each data set. Subtyping appears more precise and potentially more useful for treatment and prognoses by duly recognizing this complexity. Our approach is flexible, in the sense that if a single cluster assignment would emerge from the integrative clustering, it would appear in the TWL model, which however is able to capture deviations from it. Our analysis on the breast cancer data shows clearly that a single, unique clustering of all patients cannot account for pan-genomic sample heterogeneity and can therefore be misleading. We believe that a TWL analysis of breast cancer patients would be the best starting point to study precision treatments.

Our integrative clustering data analysis is unique in that it includes 57 women with ductal carcinoma in-situ, in addition to 313 women with invasive breast cancer. The Cancer Genome Atlas exclusively includes invasive tumors, and so our dataset and unsupervised TWL analysis is positioned to characterize the genomics of DCIS and invasive cancers in a way that is impossible in the well-analyzed TCGA samples. We leverage our integrative clustering analysis to better understand breast cancer subtypes in both DCIS and invasive breast cancer cases.

The TWL model is built to achieve robust and stable results. Because of the dimensionality of the genomic datasets, we proposed a modified likelihood with thick and even lower bounded tail values of the Gaussian model, thus coarsening the data, to achieve greater mixing in the MCMC [Hennig, 2004, Coretto and Hennig, 2016]. We experimented with simulated annealing [Kirkpatrick et al., 1983], but found convergence improved under our alternative approach.

Our Gibbs sampler was additionally modified in the burn-in period to provide a warm start to the MCMC and avoid excessive sampling of clusters close to the global mean of features in each dataset. Our ability to clearly find 5 distinct expression clusters, which align to a high degree with the known breast cancer subtypes, without specification of the number of clusters and across parallel chains and draws from the data serves as validation of our model fitting procedure on this dataset. We additionally found 2 distinct CNA clusters and 7 methylation clusters, and a relative lack of “cluster correspondence” across datasets as judged by our proposed measure.

We did not observe a high degree of pan-genomic cluster correspondence, and primarily found modest enrichment in the Her2 subtype and DCIS tumor state in those samples with greater expression-CNA cluster correspondence. There are different reasons we may not have seen significant pan-genomic cluster correspondence. According to the analysis of Sun et al. [2018], tumor purity and cell type composition confound the relationship between expression and methylation genomic data sources, and we are unable to control for these factors in our analysis. Sun et al. [2018] also finds that CNAs affect expression and methylation independently. If this is the case, one would expect weaker expression-methylation associations, and stronger associations between the expression-CNA and CNA-methylation pairs.

Myhre et al. [2013] performs a similar analysis to examine correspondence across genomic platforms, calculating Spearman correlation on gene-annotated expression, copy number, and proteomic probes both within subtypes and also marginally. Direct comparison with their results is difficult however, as they do not perform a genome-wide analysis, but examine the PI3K/Akt pathway. They do observe variation by subtype in the degree of correlation between measures on different platforms for specific genes. However, examination of all genes in the pathway does not reveal broad trends of greater correlation in one subtype over another on a specific pair of data sources. In particular, the Her2 subtype does not seem to exhibit greater expression-CNA Spearman correlation for the genes interrogated than the other subtypes.

We found that nearly all DCIS tumors inhabit the large CNA cluster identified in our analysis in a robust fashion. There were similar and perhaps more significant findings in the expression dataset, where one cluster seemed to contain most DCIS observations (see Table S13). The large CNA cluster also contained a large proportion of invasive tumors, however, and the second cluster in the CNA analysis contained almost entirely invasive tumors. The finding of a single CNA cluster for all DCIS and some IDC seems to stand in contrast to findings of Lesurf et al. [2016], who did not identify genomic loci that consistently separated the two tumor states. The difference could result from the accumulation of small differences between the two tumor states. This in turn could lead to the fairly characteristic differences one sees in the two CNA clusters in our analysis and to which other methods are not sensitive. Also unclear was why we observed so little CNA cluster homogeneity in the Basal intrinsic subtype so that most of its samples had the “unknown” cluster classification. The finding suggests that nearly all Basal samples are unlike one another and unlike any other subtype with respect to CNAs.

It may not be surprising that we observe relatively little commonality between expression and methylation clusterings. Moarii et al. [2015] finds that the correlations that do exist in breast and other cancers are in certain, tissue-specific sets of genes. While their analysis only includes invasive tumors from TCGA, genome-wide analysis therefore may very well not detect such a specific subset. We also may not expect a lot a correlation between expression clusters and those of CNAs, manifested either through posterior cross-tabulations of cluster labels from those data sources or the more global measure of between-dataset cluster correspondence. Indeed, when Curtis et al. [2012] integrates the two sources of information, they find CNAs stratify subtypes in some cases and also group subsets from disparate ones together, resulting in 10 total integrated clusters.

The volume of genomic data will only increase, and integrative clustering models have an important role to play in giving insight into underlying biology. The TWL model, because of its scalability and flexibility, can aid in this understanding.

## 5 Data Availability

Data is available within our published R package on CRAN as “twl” [R Core Team, 2017].

## Supporting information

Supplementary methods and analyses

## 6 Acknowledgements

We acknowledge useful discussions with Sylvia Richardson and Paul Kirk. Funding for this research was also provided by the Research Council of Norway, BigInsight, the Norwegian Cancer Society, and the South-Eastern Norway Regional Health Authority. We thank Dr. Maria Grazia Daidone (Fondazione IRCCS Instituto Nazionale dei Tumori, Milan, Italy) for the contribution of patient samples to the project.

