## Supplementary methods and analyses for "A Bayesian Two-Way Latent Structure Model for Genomic Data Integration Reveals Few Pan-Genomic Cluster Subtypes in a Breast Cancer Cohort"

#### S1.1 Gibbs Sampler implementation

##### S1.1.1 Tilting the log likelihood for unoccupied clusters during MCMC burn-in

We begin now to describe aspects of the Gibbs Sampler implementation of our model. There was practical use for choosing default values of the log-likelihood for unoccupied clusters or those with only 1 observation during the burn-in of the MCMC to speed convergence. The suggested priors on the cluster means and the use of global variance parameters lead to full conditionals for moves to unoccupied clusters. However such a model favors observations whose feature vectors are close to the mean feature vector across all samples to be sampled into unoccupied clusters. Samples more distant from the mean feature vector across all samples are less likely to be sampled into unoccupied clusters, and when such samples are, the prior on the cluster mean parameters will exert relatively greater influence on the resulting posterior. As a result, sampling still more distant samples into the unoccupied cluster is rendered less likely.

We calculate what the log-likelihood of the Gaussian mixture model would be if in truth samples were generated from M clusters, but we performed random cluster assignment of them. That is, they are assigned a label not corresponding to the one from which they come, but any random integer in $\{1, \ldots, M\}$, and cluster mean and variance parameters are calculated assuming that random cluster assignment. One can think of this random assignment as that which would be "observed" in the absence of any information or as a first guess in an iterative or sampling fitting procedure. Suppose the marginal variance (i.e., the vector of variances of features across all samples in a data set irrespective of cluster assignment) is $\sigma^2$ and the assumed within-cluster and between-cluster variances are $c \cdot \sigma^2$ and $(1-c) \cdot \sigma^2$ for some $c \in (0, 1)$. (One must choose $c$, and 0.5 is suggested as a conservative choice. In practice, the calculated log likelihood is relatively insensitive to choice of $c$.)

Consider $E_{f_{\mu_2}}[E_{f_{\mu_1}}[E_g[log\, h(X)|\mu_1, \mu_2]|\mu_1]]$ where $\mu_1$ and $\mu_2$ are treated as random variables and represent the mean values at which the Gaussian mixture model likelihood is evaluated (i.e., where cluster-specific mean parameters assume the random cluster assignment) and from which the observation is generated (i.e., the mean parameter of the true cluster), respectively. This is

$$\int \left( \int \left( \int log(h(x))\, g(x)\, dx \right) f_{\mu_1}(\mu_1)\, d\mu_1 \right) f_{\mu_2}(\mu_2)\, d\mu_2$$

where $h$ is a Gaussian pdf used to evaluate our samples with randomly assigned cluster labels. It is indexed by mean parameter $\mu_2$ and variance $\sigma^2$. The Gaussian density $g$ is the sample's true density, and $f_{\mu_1}$, and $f_{\mu_2}$ are also Gaussian pdfs for the means of $g$ and $h$, respectively. So informally, we are evaluating the log likelihood with h, which corresponds to the random (or "observed") cluster assignment, according to the density of the truth, g, where what determines truth is the indexing of these densities by $\mu_1$ or $\mu_2$. Additionally, $g$, $f_{\mu_1}$, and $f_{\mu_2}$ are all indexed by variance parameter $\sigma^2$ and by mean parameters $\mu_1$, 0, and 0, respectively (the 0's chosen arbitrarily since the result holds so long as these numbers are equal). So $h = h(\cdot|\, \mu_2, c \cdot \sigma^2)$, $g = g(\cdot|\, \mu_1, c \cdot \sigma^2)$, $f_{\mu_1} = f_{\mu_1}(\cdot|\, 0, (1-c) \cdot \sigma^2)$, and $f_{\mu_2} = f_{\mu_2}(\cdot|\, 0, (1-c) \cdot \sigma^2)$. We expand the expression above to

$$= \int \left( \int \left( \int -1/2 \cdot \log C - \frac{(x - \mu_2)^2}{2 \cdot c\,\sigma^2}\, g(x)\, dx \right) f_{\mu_1}(\mu_1)\, d\mu_1 \right) f_{\mu_2}(\mu_2)\, d\mu_2$$

where $C = 2\,\pi\sigma^2$, then have

$$= -1/2 \cdot \log C - 1/2(c) - \int \left( \int \frac{(\mu_1 - \mu_2)^2}{2 \cdot c\,\sigma^2}\, f_{\mu_1}(\mu_1)\, d\mu_1 \right) f_{\mu_2}(\mu_2)\, d\mu_2$$

$$= -1/2 \cdot \log C - 1/2(c) - \int \left( \int \left( \frac{\mu_1^2}{2 \cdot c\,\sigma^2} + \frac{\mu_2^2}{2 \cdot c\,\sigma^2} \right) f_{\mu_1}(\mu_1)\, d\mu_1 \right) f_{\mu_2}(\mu_2)\, d\mu_2$$

$$= -1/2 \cdot \log C - 1/2 \cdot (c + 2\,(1 - c))$$

This formula has been confirmed in simulation. We choose $c = 0.5$ though note that since $\log C$ dominates the expression, choice of $c$ is not critical. While ad hoc, we found that using this formula for the default likelihood effectively allows for the repopulation of clusters once having become unoccupied, maintains cluster sparsity, and does not favor repopulating unoccupied clusters with observations closer to the global mean. This strategy was only used during the burn-in of the MCMC so that it would not distort samples from the posterior.

##### S1.1.2 Modified densities

Sampling from the conditional posterior for $C_{ij}$ shown in Eqn. (1) will sometimes be dominated by one of the three terms of each normalized multinomial category. This is especially true for the likelihood model–the higher the dimension of the data, the more peaked one cluster likelihood score will tend to be relative to all others. And since the Gaussian model has thin tails, the corresponding log likelihood can become very small without lower bound for many clusters. This is unlike the priors on $p_j$ and $\rho_i$, which effectively have thick "tails" because of hyperparameters $\alpha$ and $\beta$ giving non-trivial probability mass to sampling even unoccupied clusters. As a result, even in settings where clusterings in other datasets are similar, the model will not tend to sample similar cluster labels across them. Instead, when sampling the posterior one will quickly find and stay near local maxima, and there will be little or no mixing of cluster assignments as the high-dimensional likelihood model renders the probability of sampling most cluster labels zero.

Since a primary feature of our model is the way in which cluster assignment information is shared across datasets and posterior assessment of across-dataset "cluster correspondence", this characteristic renders the basic model of little practical use. We can address this concern by choosing to lower bound the likelihood model, effectively censoring values that fall below the threshold to that value. One can choose the threshold as a function of the ceiling on the number of clusters (ie, $K$), and the minimum cluster label mixing desired per MCMC sample. This strategy is not without precedent and has recently been developed by Coretto and Hennig [2016] and Coretto and Hennig [2017], though it has past foundation in Hennig [2004] and Banfield and Raftery [1993]. Raftery, in particular, would envision another level of mixture for our within-dataset mixture model, that of an improper noise component.

This approach is more helpful for achieving adequate cluster label mixing than use of thick-tailed densities, which again get stuck in local maxima. Indeed, when notions of distance between cluster centers is only significant up to some threshold, use of modified densities is fitting. It is important to notice that use of such densities does not dull posterior peaks, whose functional form remains unaffected by lower bounding the tails. Our data analysis in Section 3 demonstrates that clusters still remain highly visible with use of these modified densities. One can also simply run an MCMC longer and examine statistical significance in a PSM if there is concern about slightly less visible clusters. We also experimented with simulated annealing to address the problem of the likelihood dominating posterior sampling, but found more robust results using modified densities.

#### S1.2 Comparison with Cluster of Cluster Assignments and iCluster data integration methods

To compare our TWL model with those assuming a set of common boundaries across all data types, we fit iCluster and COCA models on our data Shen et al. [2009], Curtis et al. [2012]. We fit the iCluster model with a total of 10 clusters in keeping with previously used number of subtypes in breast cancer based on multitype data (Hoadley et al. [2014]). Examination of BIC additionally suggested this was an adequate number of clusters to represent the data. Consistent with previous use of the COCA procedure, we first performed non-negative matrix factorization (NMF) on each data type and chose the number of clusters in each one suggested by our heatmaps (5, 2, and 7 for Expression, CNA, and Methylation, respectively). We then used the weighting matrices from NMF as indicator matrices for subtype and specified a total of 12 consensus clusters using the *ConsensusClusterPlus* package in R (R Core Team [2017]). Because both iCluster and COCA require all datatypes for each sample, contrary to TWL, we could only analyze 299 of our total of 370 samples.

iCluster and COCA clusterings are shown in Figures 8a, 8b, and 8c as "barcode" annotation beneath the PSM heatmaps. Colors in the barcodes map to arbitrary cluster labels and were chosen to maximize color contrast in the annotation. Patients missing at least one datatype and therefore lacking a consensus cluster are denoted with black. Since the output of both methods is a single, consensus cluster, the different barcodes under each datatype are reorderings of the consensus cluster based on sample ordering of the heatmap.

We see considerable agreement between iCluster and COCA labels and clear association with certain heatmap clusters: the enrichment in iCluster and COCA cluster labels under the methylation heatmap clusters and the Basal expression heatmap cluster (denoted by red in the "subtyp" barcode annotation) are particularly striking. However, for other expression clusters and particularly the CNA datatype as a whole, the consensus clusterings of iCluster and COCA show little consistency with datatype clusters as revealed in the heatmaps. This phenomenon likely results from the models' attempts to find consistency across datatype clusterings while there is little. For COCA in particular, its greater association with the methylation heatmap clustering may come in part from the larger number of clusters in that datatype and resultant greater influence on the consensus clustering. Ultimately there seems to be much loss of datatype-specific clustering information by fitting a single consensus cluster for each sample.

##### S1.2.1 Label switching

Label switching due to posterior symmetry in label permutations is not a concern in our model [Rodríguez and Walker, 2014]. Cluster label is never of direct importance in post-processing our posterior – we only examine pair-wise common cluster assignment of observations, generating a dataset-specific $N \times N$ indicator matrix for each iteration in the MCMC chain after convergence, called posterior similarity matrix (PSM) [Fritsch and Ickstadt, 2009]. A "1" in the $(v, t)$-th element of the indicator matrix associated with dataset j and iteration m indicates observations $v$ and $t$ in dataset $j$ have a common cluster assignment, say "12", for iteration $m$. We can additionally average over all iterations for each dataset to generate a correlation matrix and use either hierarchical clustering or graphical lasso to define definite, fixed clusters, if one wants [Bien and Tibshirani, 2011, Ward, 1963]. In practice, we have used hierarchical clustering for computational ease and subsequent flexibility of defining the number of clusters by simply "cutting" the dendrogram at different heights. We describe these post-processes more in Section 3.2.

We also recommend viewing the correlation matrix associated with each data set as a final product – the information loss associated with thresholding for definite cluster assignment can negatively impact more subtle conclusions from analysis. Ambiguity in cluster assignment made evident in the matrix is relevant information, and alignment or its lack with clusters found in other datasets can be unstable when thresholding rules are used to define object-dataset cluster membership.

### S2 Parallel analyses and additional model runs

#### S2.1 Genomic Instability Index, CNA clustering, and "core Basal"

The clustering for the CNA data type in our analysis is interpreted differently and shows a lack of a clear cluster for the Basal subtype (no dark blue square from the posterior similarity matrix above the bright red subtype annotation in Figure 8b), but, critically, *only* that subtype. This was an interesting feature of the analysis, and one that became more so once the genomic instability index (GII) was plotted underneath the samples. In Figure S3 below, we see a marked spike in GII under these Basal samples lacking a clear cluster.

This spike is a perhaps expected result statistically and consistent with biology including Curtis et al., who write in their manuscript, "We also noted that the majority of Basal-like tumours formed a stable, mostly high genomic instability subgroup...". From a statistical perspective, it is to say that, for CNAs, Basal samples are indeed genomically isolated (similar to conclusions from Expr and Methyl), though in a slightly different way as these samples are so heterogeneous among one another that our model cannot cluster them together. This is consistent with their very high GII and also points to a critical difference between interpretation when clustering with a mixture model (ie, our model when conditioned on a single data type) versus hierarchical clustering and dendrograms: those samples that dendrograms merge into larger clades very high on the tree are those that mixture models would tend to leave unclustered, which is synonymous with our "clus_unknown" label (see Figure S4, where we color code CNA clusters in subfigure a) and show height at which the sample is merged into a larger clade in subfigure b)). The black color indicates samples with the clus_unknown label, and we see that these samples are merged higher on the dendrogram than other samples. We therefore try to clarify here again our meaning of "clus_unknown" – it doesn't mean that "this sample belongs to a cluster but we don't know which one". Rather, it means "this sample is so different from everything else receiving a cluster label that we are forced to leave it without a cluster label". Whereas we have insight into samples deserving the clus_unknown label using mixture models like TWL, we would likely not have gained that insight had we simply performed hierarchical clustering.

Many of these isolated, Basal clusters in all data types additionally seem to correspond to what the literature has termed 'core Basal', a subtype in our sample which is entirely invasive and different than other, largely DCIS, Basal samples (Blows et al. [2010], Cheang et al. [2008]). Curtis also describes the notion of core Basal without naming it as such. The Basal cluster particularly clear in the expression clustering for our TWL model exhibits core Basal characteristics. If one color codes these Basal samples in red versus other Basal samples in our cohort in blue and plots all against correlation to Basal and Luminal A (the "opposite" subtype) PAM50 centroids, one sees clear separation between the two kinds of samples (see Figure S5). Irrespective of the TWL clusterings, if one simply considers all subtypes, but stratifies by DCIS/Inv, similar genomic isolation of Basal is seen among the invasive samples (data not shown). If one makes the plot among DCIS samples, however, one sees no such separation. This is all to say that a notion of core Basal exists in the literature and describes samples with highly Basal-like characteristics, these samples tend to be invasive, and the expression clustering in the TWL run was sensitive to finding these samples without any a priori knownledge of them.

#### S2.2 Clustering with promoter-region methylation probes

A unique feature of our study and one that distinguishes it from TCGA is its inclusion and focus on DCIS samples. As written in the main text, we collapsed certain methylation probes because doing so showed greater success in distinguishing DCIS/Inv tumors using penalized regression models based on internal research. A goal for this project was to see if unsupervised clustering could distinguish DCIS and invasive samples and so we used this same summarization method for the samples we clustered.

While probe localization is very important in, for example, determining methylation's relationship with expression, among other things, this is not to say that CpGs in gene bodies versus promoters do or do not contain redundant information statistically. It is possible and a different question whether the profiling derived from collapsed probes is very similar to that derived from, for example, promoter regions, despite promoter region methylation having specific effects on aspects of biology.

We have investigated this question in the analyses below, whether CpG annotation does or does not lead to different clusterings of subjects. First, we establish that our original methylation clustering was detecting structure in the collapsed probes. See Figure S6 for a dendrogram whose leaves are color coded according to our model-defined methylation clusters. Common colors of adjacent leaves makes clear that our model is detecting structure in the data we analyzed. Figure S1c similarly shows the relationship between underlying patterns in the methylation data and the clustering in our analysis. Next, we performed hierarchical clustering and tanglegram analyses (two dengrograms mapped against one another for comparison) with stratified datasets restricted to gene bodies, promoter regions, and where probes are located with respect to CpG islands. See Figure S7a for an example comparison between structures in the original, collapsed dataset with uncollapsed probes. For comparison, we include a tanglegram of the same datasets but generated under the null hypothesis of no common structure by randomly permuting sample labels (see Figure S7b). Tanglegrams for datasets restricted to the different probe annotations look similar. These figures show that there are many common structures between the collapsed and uncollapsed datasets.

In one of the two additional TWL analyses we ran in this Supplement, our methylation dataset contained only probes in promoter regions. For that run, we indeed see some changes in the methylation clustering in Figure S2 where in each subfigure samples are ordered according to clusterings in the main text (or see top portions of each subfigure in Figure S1) for easy comparison. However, some features of the original clustering are clearly maintained: clusters from that clustering are split between 3 clusters in the new clustering in only one case and split between 2 or maintained in all other cases. This analysis run also used different hyperparameters. Related to the change in beta hyperparameter, one sees slightly lighter shades of blue in the expression and CNA clusterings of Figure S2 compared to clusterings in the main text.

Ordering the new, only promoter region, methylation data according to its own clustering and attaching PAM50 subtype annotation gives Figure S8. So there seem to be a similar number of clusters as those with the uncollapsed data, and those clusters have relatively high correlation with the previous set of methylation clusters as subfigure c) of Figure S2 shows. The fact that there are slightly fewer samples within a definite cluster (fewer samples with a blue block above them in Figure S8 interestingly enriched for the Her2 subtype) in this clustering comes from the increase in the beta hyperparameter.

#### S2.3 Comparison with past integrative breast cancer genomics

We first focus on the similarities between our results and those of Curtis et al and those of the TCGA consortium. The Basal subtype is nearly perfectly restricted to a single cluster in the expression data type clustering resulting from our model (Figure 8a of the main document, bright red of the subtype annotation 'barcode' lining up with the square cluster above it. Table S4 in this supplementary material also shows the perfect specificity of this cluster and very good sensitivity). The smallest cluster in the methylation clustering (Figure 8c of the manuscript) additionally has perfect specificity for the Basal subtype (again the red annotation below the small, blue square of the posterior similarity matrix). There are many Basal samples in an adjacent, related cluster in the methylation clustering and that cluster is highly enriched for this subtype. We already explained above how Basal is genomically isolated in a different way in the CNA clustering.

There are reasons our analyses are not directly comparable with Curtis et al and the TCGA consortium. We used features sampled genome-wide, while Curtis, in choosing copy number features, seemed to filter them according to those most correlated with expression (p. 25 of their supplement). In this sense their feature selection better aligned with assumptions of iCluster, that common clusterings can be found across data types and may have significantly affected copy number clusterings for that data type. Additionally, iCluster itself can select those features more consistent with the common clustering assumption. The TCGA consortium seems to use non-negative matrix factorization for cluster assignment. TCGA filters their methylation probes based on standard deviation, which we did not do. Our collapsed methylation probes (we collapsed because it seemed to better distinguish DCIS/Inv) came from 450K arrays whose probes are a fairly representative sample of TSS's, gene bodies, and position relative to CpGs. While TCGA uses 450K arrays for some samples, they seem to filter down to those found in 27K arrays, which are nearly exclusively found in promoter regions. Lastly, our data includes DCIS samples which could affect clusterings and which neither the Curtis nor TCGA analyses do. So there are a variety of sources of variation between the two sets of analyses.

### S3 The general two-way latent structure model

Here we present our model in the important situation where samples have measurements only in some of the data sets. $A_{ij}$ is the subject id associated with row $i$ in dataset $j$. It is only annotation, or metadata, and serves as the index to which we apply multinomial models across datasets, within sample. If all data sources have identical sets of sample ids, the notion of $A_{ij}$ is unnecessary and could be reduced to $i$ since rows in each data set could be associated with the same sample if ordered properly. It is because we seek to fit our model in settings where some subjects only have a subset of data sources that we introduce $A_{ij}$ notation. We assume $A_{ij} \in \{id_1, \ldots, id_n, \ldots, id_N\}$, the superset of ids over all datasets

Let $j$ be the dataset index, and we assume $j \in \{1, \ldots, J\}$, for a total of $J$ data sources.

$Y_{i,j}$ is a vector of features for id $A_{ij}$ and data set j and of dimension $d_j$. Not all sample-dataset pairs $(A_{ij}, j)$ need exist if a certain data source is not available for sample $id_n$. Assume that the cardinality of the ids in data set $j$ is $N_j$. So $\forall\, j$, $N_j \leq N$, $N$ again the size of the superset of ids. Dataset $j$ therefore consists of $N_j$ observation vectors $Y_{A_{ij},j}$ of length $d_j$.

Let $\mathbf{Y}$ be the set of all observation vectors $\{Y_{i,j}\}$ for all valid pairs of $(i, j)$

We seek to sample from the posterior:

$$P(\boldsymbol{\mu}, \boldsymbol{\tau}, \mathbf{p}, \boldsymbol{\rho}, \mathbf{R}, \mathbf{C} \,|\, \mathbf{Y}, \mathbf{C} = \mathbf{R})$$

where we now explain $\boldsymbol{\tau},\ \mathbf{p},\ \boldsymbol{\rho},\ \mathbf{R},\ \mathbf{C}$

Column (ie, data source) clusterings are defined with the random vector

$$\mathbf{C} \equiv (C_{1,1}, \ldots, C_{i,j}, \ldots, C_{N,J})$$

where $J$ again is the total number of data sets, and

$$C_{(i,j)} \sim Multinom(p_j^{(1)}, p_j^{(2)} \ldots, p_j^{(K)}) \;\; \forall \;\text{valid}\; (i, j)$$

where $K$ is the fixed upper bound on the number of clusters. The interpretation of $p_j^{(k)}$ is the probability of a draw of cluster label $k$ in dataset $j$. So the $1, 2, \ldots, K$ in parentheses here are superscripts, not exponents, and similarly for the model on the $R_{i,j}$'s below.

We emphasize that though $C$ is subscripted by $i$ and $j$, the parameters are only subscripted by $j$. Therefore, all observations within dataset $j$ draw from this dataset-specific $j^{th}$ multinomial model. We can loosely think of the cluster models on the $j$'s as models on the *Columns* of the aggregated datasets if we arranged them next to one another.

Row (ie, sample id) clusterings are defined with the random vector

$$\mathbf{R} \equiv (R_{1,1}, \ldots, R_{i,j}, \ldots, R_{N,J})$$

Assuming that $A_{ij} = id_n$, we have

$$R_{(i,j)} \sim Multinom(\rho_{id_n}^{(1)}, \rho_{id_n}^{(2)} \ldots, \rho_{id_n}^{(K)}) \;\; \forall \;\text{valid}\; (i, j)$$

with $K$ the upper bound on the number of clusters. Note that though $R$ is subscripted by $i$ and $j$, the parameters are only subsripted by $id_n$, subject id annotation. We can loosely think of the cluster models on the $id_n$'s as models on the *Rows* of the aggregated data sources if we arranged them next to one another.

The model for $\mathbf{p_j}$ is

$$\mathbf{p_j} \sim Dirichlet(\beta_1, \ldots, \beta_K) \;\; \forall \;\; j$$

And for $\boldsymbol{\rho}_{id_n}$,

$$\boldsymbol{\rho}_{id_n} \sim Dirichlet(\alpha_1, \ldots, \alpha_K) \;\; \forall \;\; id_n$$

with hyperparameters $\alpha$ and $\beta$ constant across $id_n$'s and $j$'s, respectively. $\boldsymbol{\alpha}$ should likely be chosen as a function of the average number of data sets per id (or simply number of data sets if all ids are present in all data sets and unique within them), and $\boldsymbol{\beta}$ as a function of the average number unique ids within each data set (or simply number of unique ids if again all ids are present in all data sets and unique within them). As a result, and as follows, we have

$$\alpha \equiv \alpha_1 = \alpha_2 = \cdots = \alpha_K$$

and

$$\beta \equiv \beta_1 = \beta_2 = \cdots = \beta_K$$

The parameters of the normal densities are defined with:

$$\boldsymbol{\mu} \equiv (\mu_{1,1} \dots \mu_{K,1}, \dots\dots, \mu_{1,j} \dots \mu_{K,j}, \dots\dots, \mu_{1,J} \dots \mu_{K,J})$$

where $\mu_{k,j}$ is the mean vector and of dimension $d_j$, $d_j$ again the number of features in data source $j$.

$$\boldsymbol{\tau} \equiv (\tau_{1,1} \dots \tau_{K,1}, \dots\dots, \tau_{1,j} \dots \tau_{K,j}, \dots\dots, \tau_{1,J} \dots \tau_{K,J})$$

where $\tau_{k,j}$ is the precision vector and of dimension $d_j$. Independence is assumed for components of vector $Y_{i,j}$ and as a result information of its precision matrix is contained in a $d_j$ dimensional vecor. The assumption increases computation efficiency, and filtering highly correlated features makes the assumption reasonable.

$\mathbf{C} = \mathbf{R}$ is shorthand notation for the event $\cup_{\{i,j\}} C_{i,j} = R_{i,j}$; i.e., for all valid pairs of $(i, j)$, $C_{i,j} = R_{i,j}$ is true.

We factorize the posterior to the following and consider each factor in turn in the next sections.

$$P(\boldsymbol{\mu}, \boldsymbol{\tau}, \mathbf{p}, \boldsymbol{\rho}, \mathbf{R}, \mathbf{C} \,|\, \mathbf{Y}, \mathbf{C} = \mathbf{R}) \propto$$

$$P(\mathbf{Y} \mid \boldsymbol{\mu}, \boldsymbol{\tau}, \mathbf{p}, \boldsymbol{\rho}, \mathbf{R}, \mathbf{C}, \mathbf{C} = \mathbf{R}) \cdot P(\boldsymbol{\mu}, \boldsymbol{\tau} | \mathbf{p}, \boldsymbol{\rho}, \mathbf{R}, \mathbf{C}, \mathbf{C} = \mathbf{R})$$

$$\cdot P(\mathbf{C}, \mathbf{R} \mid \mathbf{p}, \boldsymbol{\rho}, \mathbf{C} = \mathbf{R}) \cdot P(\mathbf{p}, \boldsymbol{\rho} \,|\, \mathbf{C} = \mathbf{R})$$

#### S3.1 Considering Y

In the expressions below we have define $k^* \equiv C_{i,j}$ and use $k^*$ for cleaner notation to avoid double subscripting. It functions as shorthand for subscripting according to the $C_{i,j}$ cluster label. We use $\{i, j\}$ to denote the set of valid $(i, j)$ pairs.

$$P(\mathbf{Y} \mid \boldsymbol{\mu}, \boldsymbol{\tau}, \mathbf{p}, \boldsymbol{\rho}, \mathbf{R}, \mathbf{C}, \mathbf{C} = \mathbf{R})$$

$$= \prod_{\{i,j\}} P(Y_{i,j} \mid \boldsymbol{\mu}, \boldsymbol{\tau}, \mathbf{p}, \boldsymbol{\rho}, \mathbf{R}, \mathbf{C}, \mathbf{C} = \mathbf{R})$$

$$= \prod_{\{i,j\}} P(Y_{i,j} \mid \mu_{k^*,j}, \tau_{k^*,j}, C_{i,j}, R_{i,j}, C_{i,j} = R_{i,j}) = \prod_{\{i,j\}} P(Y_{i,j} \mid \mu_{k^*,j}, \tau_{k^*,j}, C_{i,j})$$

$$= \prod_{\{i,j\}} P(Y_{i,j} \mid \mu_{k^*,j}, \tau_{k^*,j}) = \prod_{\{i,j\}} N(Y_{i,j} \mid \mu_{k^*,j}, \tau_{k^*,j})$$

where $N(\cdot)$ is used to denote the Gaussian density here and below. Each $(i, j)$ term in this last expression can be written as $\prod_{k=1}^{K} N(Y_{i,j} \mid \mu_{k,j}, \tau_{k,j})^{I(k=C_{i,j})}$. For the purpose of combining terms when calculating posteriors on the $C_{i,j}$'s, we write the likelihood as

$$= \prod_{\{i,j\}} \prod_{k=1}^{K} N(Y_{i,j} \mid \mu_{k,j}, \tau_{k,j})^{I(k=C_{i,j})}$$

Notice we arbitrarily dropped the $R_{i,j}$ because it contains redundant information on $C_{i,j}$. However, notationally the choice is not entirely arbitrary because $C_{i,j}$ can be considered in some sense "more primary" for notation since $\boldsymbol{\mu}$ and $\boldsymbol{\tau}$ posteriors are sampled based only on within dataset (or loosely, column) cluster assignment. This is also evident in our choice of $\boldsymbol{\mu}$ and $\boldsymbol{\tau}$ subscripts.

#### S3.2 Considering C, R

Now consider

$$P(\mathbf{C}, \mathbf{R} \mid \mathbf{p}, \boldsymbol{\rho}, \mathbf{C} = \mathbf{R})$$

$$= \prod_{\{i,j\}} P(C_{i,j}, R_{i,j} | \; \mathbf{p}, \boldsymbol{\rho}, \mathbf{C} = \mathbf{R}) = \prod_{\{i,j\}} P(C_{i,j} | \; \mathbf{p}, \boldsymbol{\rho}, \mathbf{C} = \mathbf{R})$$

$$= \prod_{\{i,j\}} \left( p_j^{(C_{i,j})} \cdot \rho_{A_{ij}}^{(C_{i,j})} / \Big( \sum_k p_j^{(k)} \cdot \rho_{A_{ij}}^{(k)} \Big) \right)$$

where again we drop $R_{i.j}$ to emphasize that it contains redundant information. The $C_{i,j}$'s and $k$'s in this expression are superscripts for the cluster label and not exponents.

#### S3.3 Considering $\boldsymbol{\mu}, \boldsymbol{\tau}, \boldsymbol{\psi}$

We now examine the term

$$P(\boldsymbol{\mu}, \boldsymbol{\tau} \,|\, \mathbf{p}, \boldsymbol{\rho}, \mathbf{R}, \mathbf{C}, \mathbf{C} = \mathbf{R})$$

$$= \prod_k \prod_j P(\mu_{k,j}, \tau_{k,j} \,|\, \mathbf{p}, \boldsymbol{\rho}, \mathbf{R}, \mathbf{C}, \mathbf{C} = \mathbf{R})$$

$$= \prod_k \prod_j N(\mu_{k,j} \,|\, \hat{\tau}_{k,j}, \mu_{0,j}, \psi_{0,j})$$

$$= \prod_k \prod_j N(\mu_{k,j} \,|\, \hat{\tau}_{k,j}, \mu_{0.j}, \psi_{0,j})$$

The mean and precision of $\mu_{k,j}$ are $\mu_{0,j}$ and $\psi_{0.j}$, respectively, where we set $\mu_{0,j}$ to the global mean vector of $Y_{\cdot,j}$, which is data set $j$. We set the corresponding precision parameter

$$\psi_{0,j} = \left( \frac{1}{2} \cdot \frac{\sigma_{G,j}^2}{n/30} \cdot 15 \right)^{-1}$$

for all $j$, where $\sigma^2_{G,j}$ is a vector of variances of the features of $Y_{\cdot,j}$. Setting $\psi_{0,j}$ in this way makes the assumption that half of global variation in data set j, $\sigma^2_{G,j}$, is attributable within-cluster, that the size of clusters will be 1/30 of the total sample size, $n$, and that there should be 1/15 cluster mean shrinkage to the global mean under such circumstances. If the analyst fits the model with an upper bound of more than 30 clusters, this will result in greater mean shrinkage earlier in the MCMC chain since the chain begins with random cluster assignment. If in truth there are fewer than 30 clusters in the data, will result in less shrinkage of the respective cluster means once convergence has occurred.

Such a prior also serves a practical purpose: without it, small clusters, especially those with one observation, will be resistant to becoming unoccupied since the mean parameter, $\mu_{k,j}$, will tend to be close or identical to the corresponding observation(s) and thus inflate the likelihood. This is more true if additionally cluster variance parameters, $\tau_{k,j}$'s, are cluster-specific, as estimated normal densities will diverge to infinity. By shrinking $\mu_{k,j}$ to a global mean, especially when the cluster size is smaller, the posterior of

$$P(\mathbf{C}, \mathbf{R} \mid \mathbf{p}, \boldsymbol{\rho}, \mathbf{C} = \mathbf{R}) = P(\mathbf{C} \mid \mathbf{p}, \boldsymbol{\rho}, \mathbf{C} = \mathbf{R})$$

will exhibit greater cluster sparsity.

Assuming a common variance parameter across clusters similarly keeps likelihoods from diverging and allows for more efficient estimation in the parameter. The parameter is estimated wth maximum likelihood. While the assumption of common cluster variance across clusters is unlikely to hold exactly, new clusters will form when a single one is too heterogeneous to be consistent with the common variance parameter.

Combining these priors with the likelihood given above, we can calculate posteriors for $\boldsymbol{\mu}$ and estimate $\tau_{1,j} = \tau_{2,j} = \cdots = \tau_{k,j} = \cdots = \tau_{K,j} \;\forall\; k$ via maximum likelihood, which we will denote $\hat{\boldsymbol{\tau}}_{\cdot,\mathbf{j}}$. So $\hat{\boldsymbol{\tau}}_{\cdot,\mathbf{j}}$ is a vector of length the number of features of data set $j$ invariant to cluster label, and which is specific to each data set $j$.

Because of conjugacy relationships, the posterior for $\mu_{k,j}$ is then

$$P(\mu_{k,j} \,\big|\, \mathbf{Y}_{\cdot,\mathbf{j}}, \boldsymbol{\mu}_{0,j}, \hat{\boldsymbol{\tau}}_{\cdot,\mathbf{j}}, \boldsymbol{\psi}_{\mathbf{0},\mathbf{j}}) = N\left(\mu_{k,j} \,\Big|\, \Big[n\,\hat{\tau}_{\cdot,j}\cdot\bar{\mathbf{Y}}_{\cdot,\mathbf{j}} + \psi_{0,j}\cdot\mu_{0,j}\Big]\cdot\Big[\psi_{0,j} + n\,\hat{\tau}_{\cdot,j}\Big]^{-1}, \Big[\psi_{0,j} + n\,\hat{\tau}_{\cdot,j}\Big]^{-1}\right)$$

parametrized as the mean and variance, not precision. The mean is seen to be a weighted average of the sample mean of the cluster and the prior, according to the respective precision parameters. Likewise, the variance is the inverse of the sum of the precision parameters. Both formula are familiar posterior parameters when using normal-normal conjugacy with variance $(1/\tau)$ considered fixed.

#### S3.4 Considering $\boldsymbol{p}, \boldsymbol{\rho}$

Lastly, we consider

$$P(\mathbf{p}, \boldsymbol{\rho} \,|\, \mathbf{C} = \mathbf{R})$$

$$= \prod_i \prod_j P(p_j, \rho_{A_{ij}} \,|\, \mathbf{C} = \mathbf{R}) \propto P(\mathbf{C} = \mathbf{R} \,|\, p_j, \rho_{A_{ij}}) \cdot \prod_i \prod_j P(p_j, \rho_{A_{ij}})$$

$$= \Big(\prod_{\{i,j\}} \sum_k p_j^{(k)} \cdot \rho_{A_{ij}}^{(k)}\Big) \cdot \Big(\prod_i \prod_j (p_j^{(1)} p_j^{(2)} p_j^{(3)} \dots p_j^{(K)})^{\beta} \cdot (\rho_{A_{ij}}^{(1)} \rho_{A_{ij}}^{(2)} \dots \rho_{A_{ij}}^{(K)})^{\alpha}\Big)$$

where $1, 2, \dots, k, \dots, K$ denotes superscripts, not exponents, and we make use of identical hyperparameters across the prior cluster probabilities of $\alpha$ and $\beta$ by taking each one as an exponent outside the respective parentheses.

We can collect terms for

$$P(\mathbf{C}, \mathbf{R} \mid \mathbf{p}, \boldsymbol{\rho}, \mathbf{C} = \mathbf{R}) \cdot P(\mathbf{p}, \boldsymbol{\rho} \mid \mathbf{C} = \mathbf{R})$$

and obtain

$$\propto \Big(\prod_{\{i,j\}} p_j^{(C_{i,j})} \cdot \rho_{A_{ij}}^{(C_{i,j})} / \Big(\sum_k p_j^{(k)} \cdot \rho_{A_{ij}}^{(k)}\Big)\Big) \cdot$$

$$\Big(\prod_{\{i,j\}} \sum_k p_j^{(k)} \cdot \rho_{A_{ij}}^{(k)}\Big) \cdot \Big(\prod_i \prod_j (p_j^{(1)} p_j^{(2)} p_j^{(3)} \dots p_j^{(K)})^{\beta} \cdot (\rho_{A_{ij}}^{(1)} \rho_{A_{ij}}^{(2)} \dots \rho_{A_{ij}}^{(K)})^{\alpha}\Big)$$

$$= \Big(\prod_{\{i,j\}} p_j^{(C_{i,j})} \cdot \rho_{A_{ij}}^{(C_{i,j})}\Big) \cdot \Big(\prod_i \prod_j (p_j^{(1)} p_j^{(2)} p_j^{(3)} \dots p_j^{(K)})^{\beta} \cdot (\rho_{A_{ij}}^{(1)} \rho_{A_{ij}}^{(2)} \dots \rho_{A_{ij}}^{(K)})^{\alpha}\Big)$$

$$= \prod_j \prod_{i|j} \Big((p_j^{(1)})^{\beta + \sum_{i|j} I(C_{i,j}=1)} \cdot (p_j^{(2)})^{\beta + \sum_{i|j} I(C_{i,j}=2)} \dots (p_j^{(K)})^{\beta + \sum_{i|j} I(C_{i,j}=K)}\Big) \cdot$$

$$\Big((\rho_{A_{ij}}^{(1)})^{\alpha + \sum_{j|i} I(R_{i,j}=1)} \cdot (\rho_{A_{ij}}^{(2)})^{\alpha + \sum_{j|i} I(R_{i,j}=2)} \dots (\rho_{A_{ij}}^{(K)})^{\alpha + \sum_{j|i} I(R_{i,j}=K)}\Big)$$

where we use the notation $i \mid j$ and $j \mid i$ to denote valid values of $i$ and $j$ within strata of $j$ and $i$, respectively.

#### S3.5 Cluster posteriors

Using $C_{i,j} = R_{i,j}$ in the previous expression and incorporating it with the likelihood, we have

$$= \prod_j \prod_{i|j} \Big((p_j^{(1)})^{\beta + \sum_{i|j} I(C_{i,j}=1)} \cdot (p_j^{(2)})^{\beta + \sum_{i|j} I(C_{i,j}=2)} \dots (p_j^{(K)})^{\beta + \sum_{i|j} I(C_{i,j}=K)}\Big) \cdot$$

$$\Big((\rho_{A_{ij}}^{(1)})^{\alpha + \sum_{j|i} I(C_{i,j}=1)} \cdot (\rho_{A_{ij}}^{(2)})^{\alpha + \sum_{j|i} I(C_{i,j}=2)} \dots (\rho_{A_{ij}}^{(K)})^{\alpha + \sum_{j|i} I(C_{i,j}=K)}\Big)$$

$$\Big(N(Y_{i,j} \mid \mu_{k,j}, \tau_{k,j})^{I(C_{i,j}=1)} \cdot N(Y_{i,j} \mid \mu_{k,j}, \tau_{k,j})^{I(C_{i,j}=2)} \dots N(Y_{i,j} \mid \mu_{k,j}, \tau_{k,j})^{I(C_{i,j}=K)}$$

which admits the kernel of a multinomial probability mass function in $C_{i,j}$.

### S4 Supplementary Figures

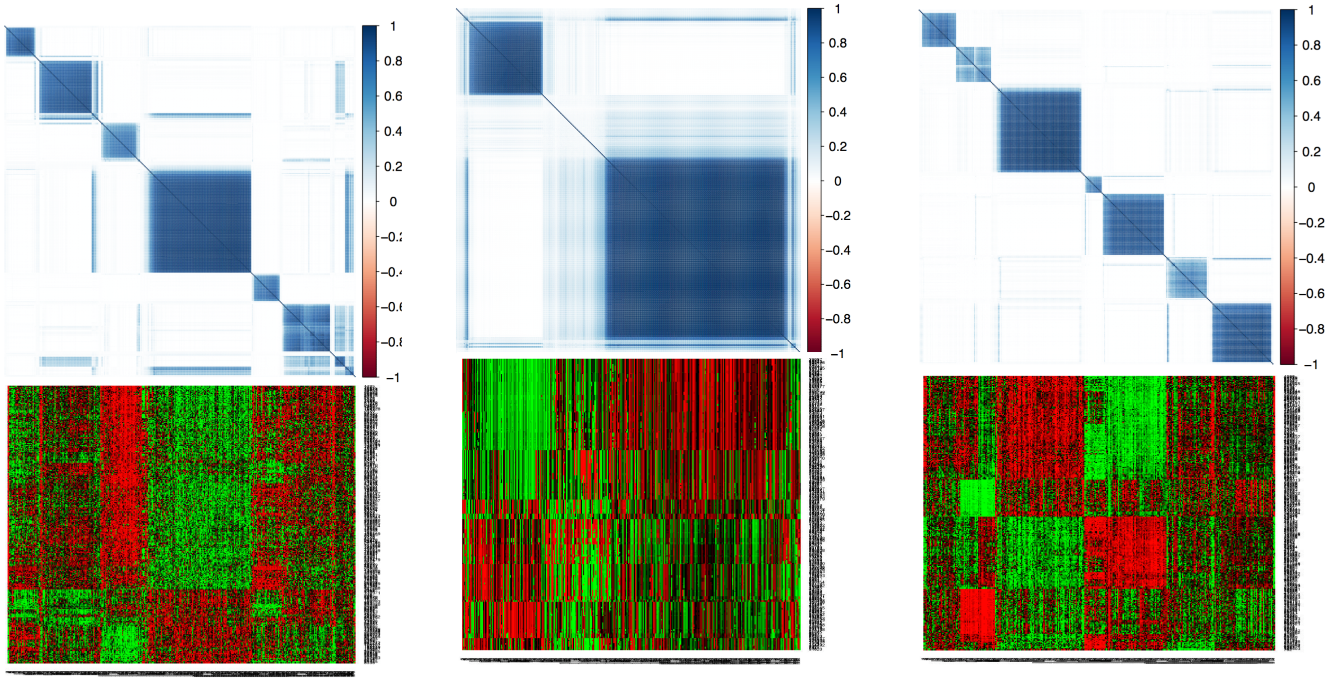


*(a) Expression* *(b) CNA* *(c) Methylation*

*Figure S1: Posterior similarity matrix clusterings from the analysis in the original manuscript with data matrix heatplots underneath. Samples of the heatplots are ordered according to those of the posterior similarity matrices and are filtered for the 15% most differentially represented between clusters for clarity. The heatplots make clear the underlying different feature patterns driving the clusterings found by our model.*

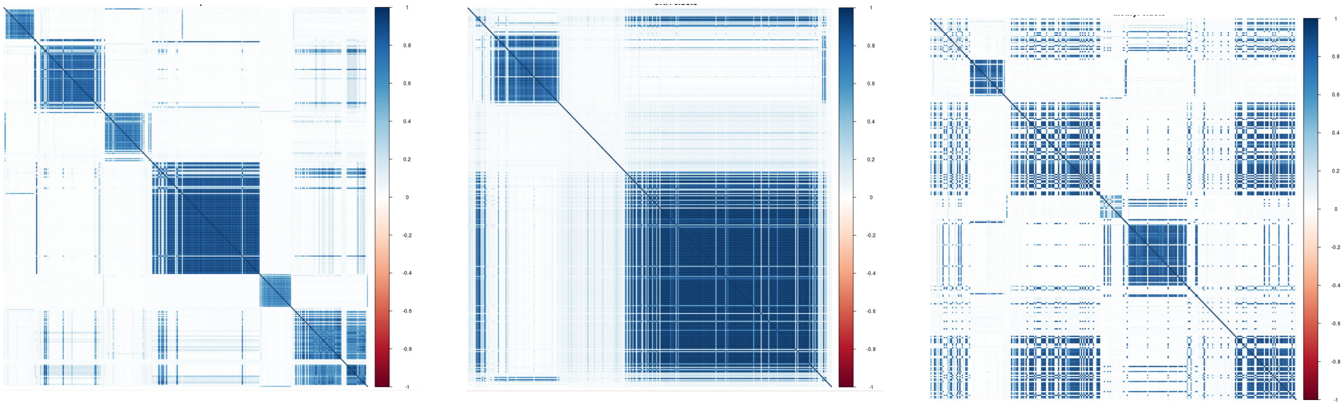

*(a) Expression dataset posterior similarity matrix* *(b) CNA dataset posterior similarity matrix* *(c) Methylation dataset posterior similarity matrix*

*Figure S2: Posterior similarity matrices for one of the two additional TWL model runs for this Supplement. We use hyperparameters alpha=0.9, beta=15, restrict our unoccupied cluster strategy to the MCMC burn-in phase, and slightly flatten the prior on $\mu_{0,j}$. We also use an entirely different dataset for methylation, 3700 uncollapsed probes from promoter regions. PSMs are ordered according to those used in the original clusterings, shown in Figure S1 for comparison. Despite these changes, clusterings in the original analysis are robust, with primary differences in the methylation dataset due to use of the alternative dataset.*

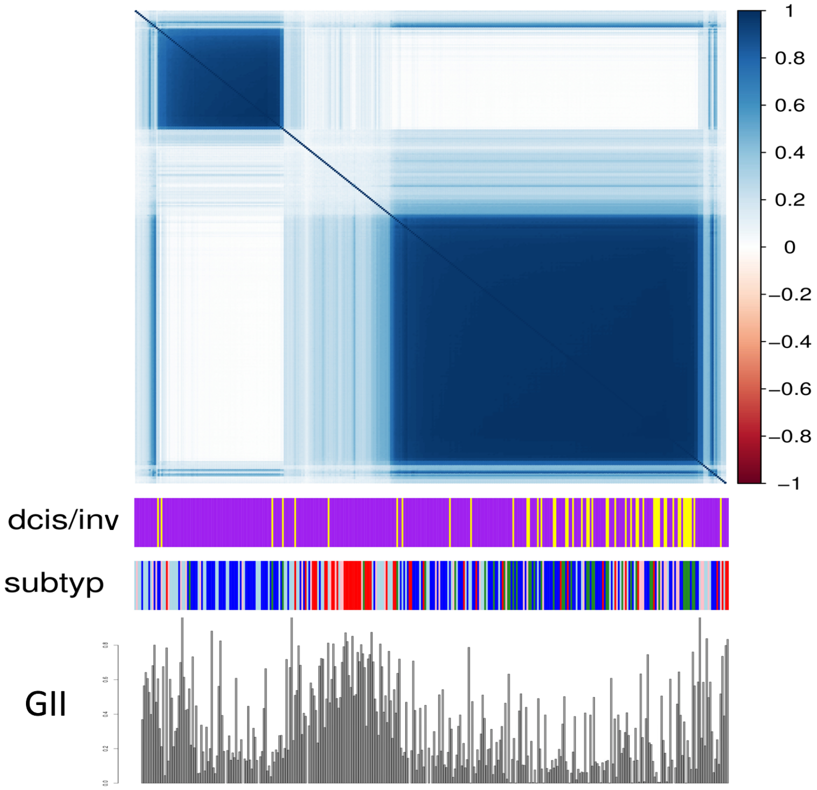


*Figure S3: TWL posterior similarity matrix from the CNA clustering in the original manuscript with subtype and DCIS annotation, plotted above the samples' genomic instability index. The Basal cluster in red has significantly higher GII and resultantly no cluster in the PSM.*

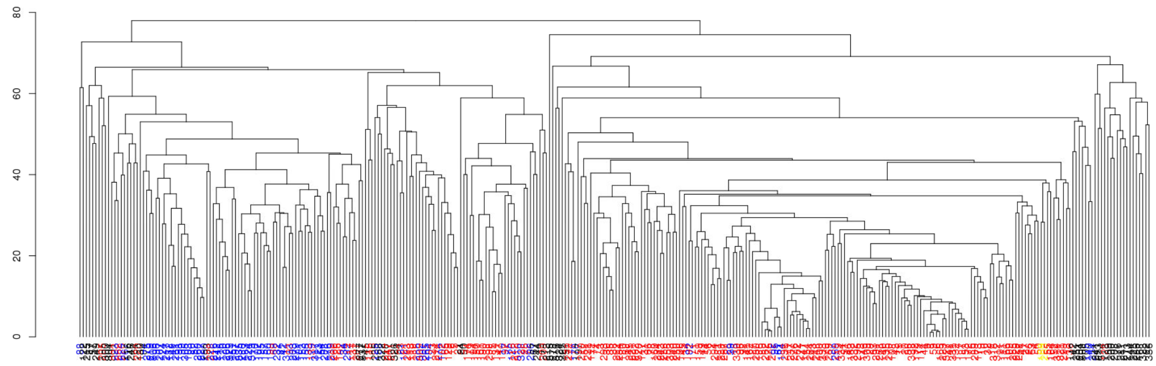

*(a) Dendrogram of CNA data, colored according to cluster from the TWL model. Black is used to denote the unknown cluster and corresponds to black points in subfigure b) below, those samples that are joined to larger clades high in the dendrogram. This feature indicates their distinct copy number profile.*

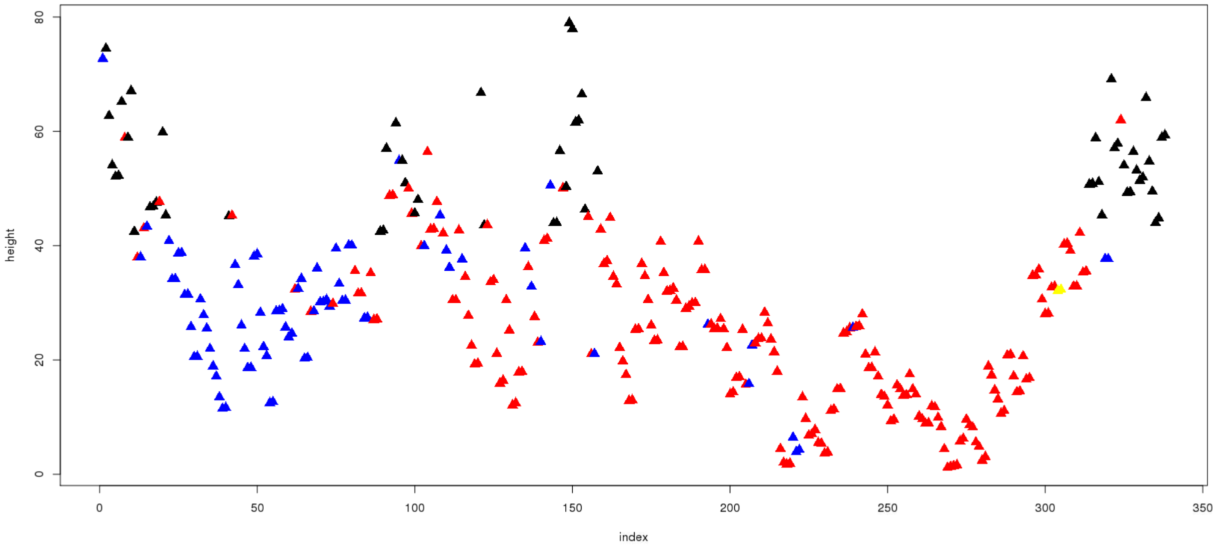


*(b) Color coded TWL clusters analyzed with hierarchical clustering and plotted according to the height at which each sample is merged into a larger clade. The higher point at which the sample is merged, the more distinct it is from the other samples.*

*Figure S4: TWL model-defined color coded leaves of a dendrogram of the CNA data in (a), aligned with height at which a leave joins a large clade in (b). "Clus_unknown" is color coded black and is exclusively found among samples being joined at higher heights in the dendrogram, indicating how different these samples' features are from other samples.*

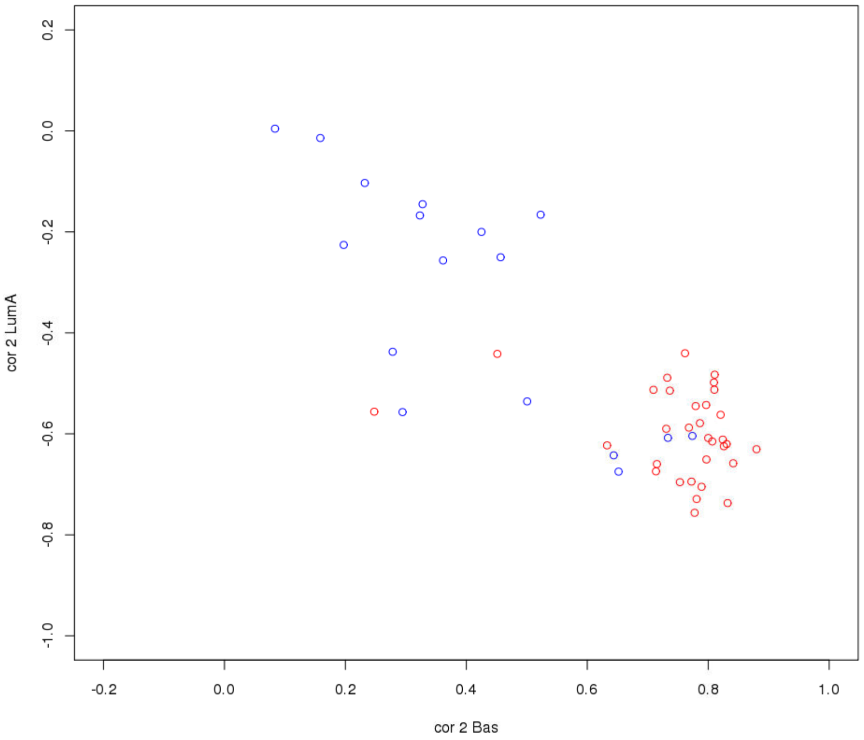


*Figure S5: Among Basal samples clustered with the TWL model, those in the clear Basal cluster in the expression clustering (color coded red here), exhibit core Basal characteristics, whereas the Basal samples in the complement (color coded blue here), do not exhibit core Basal characteristics according to their correlation with Basal and Luminal A centroids. The posterior of the TWL model reported in the original manuscript independently separates these two kinds of samples.*

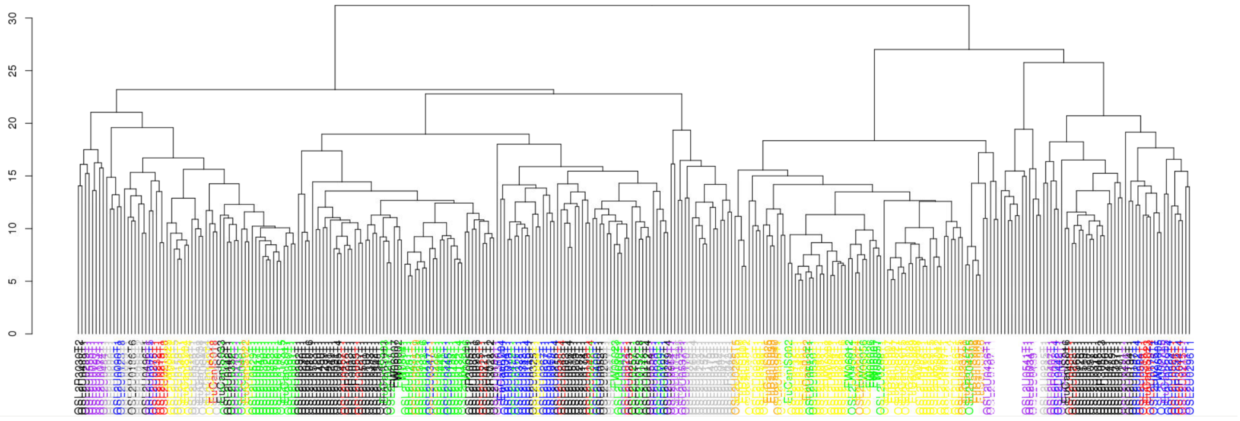


*Figure S6: Methylation data used in the original manuscript analyzed with hierarchical clustering and samples color coded according to the labels from the TWL model. Substantial clustering of the colors using the dendrogram's ordering indicates consistency between the TWL model and well-tested hierarchical clustering*

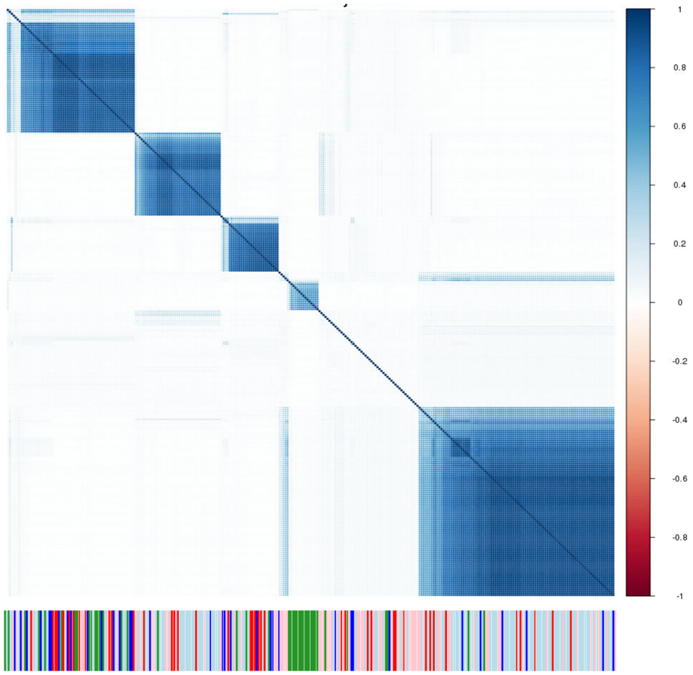


*Figure S8: Posterior similarity matrix for the TWL clustering of a methylation dataset of approximately 3700 probes from promoter regions along with intrinsic subtype annotation (red–Basal, dark blue–Lum A, sky blue–Lum B, green–normal-like, pink–Her2). This is the same matrix as that shown in Figure S2c, but ordered according to samples' own similarity, rather than ordering according to the methylation analysis in the original manuscript. The relative lack of clustering of samples immediately to the right of center of the matrix results from using a large beta hyperparameter and indicates methylation heterogeneity among these samples. These samples seem to be enriched for the Her2 intrinsic subtype, hypothetically indicating heterogeneity in the methylation profiles of that subtype.*

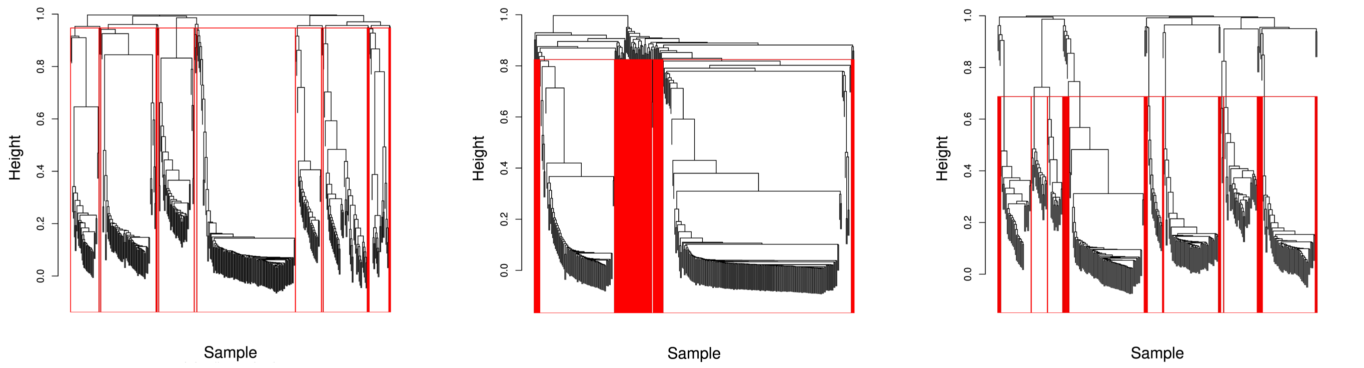


*(a) Expression dataset* *(b) CNA dataset* *(c) Methylation dataset*

*Figure S9: Post-processed hierarchical clustering for our breast cancer analysis. Hierarchical clusters generally align well with the clearly identifiable clusters in Figure 8*

### S5 METABRIC Figures

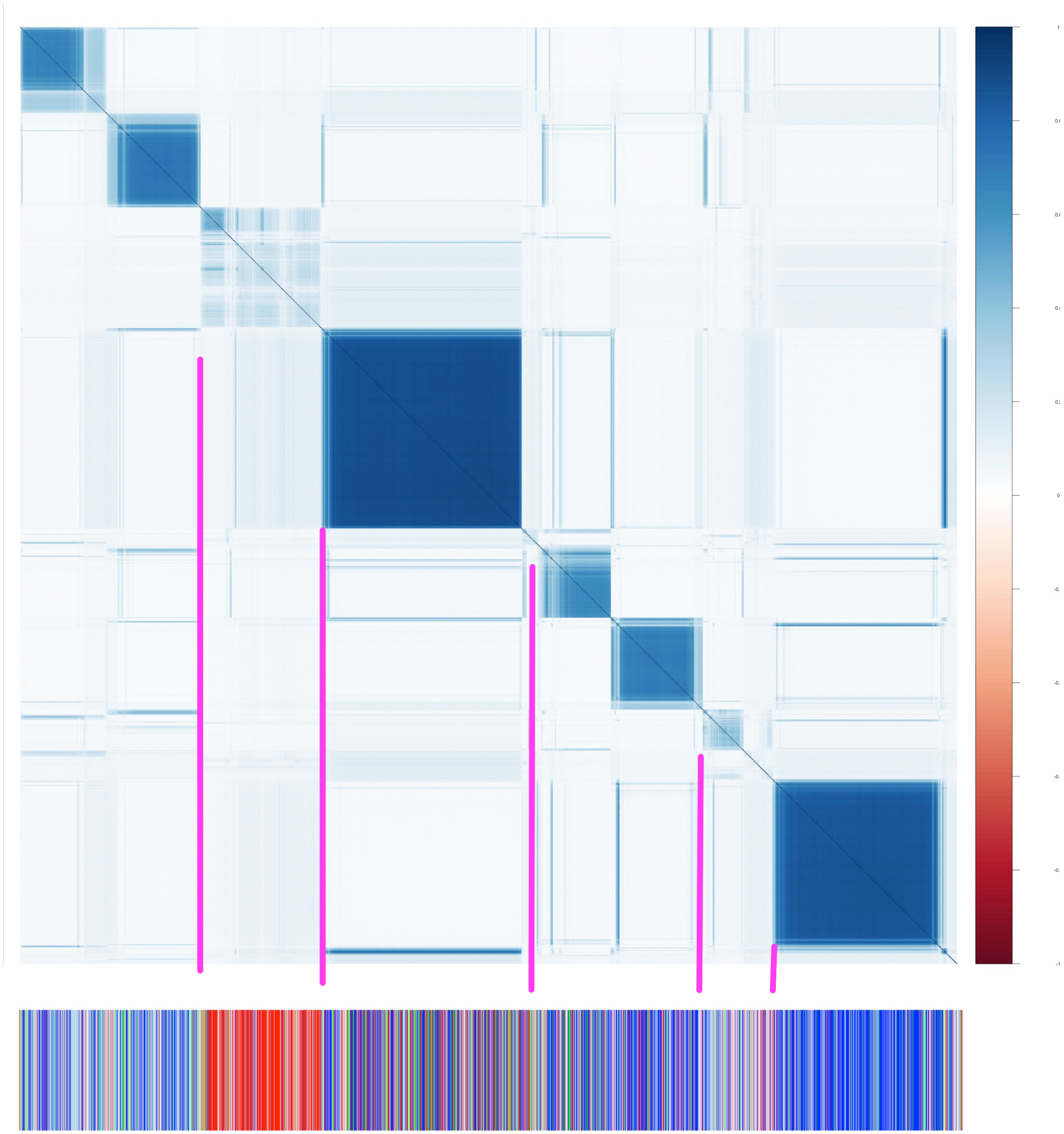

*Figure S10: CNA clustering posterior similarity matrix with subtype annotation of all 2173 subjects from Metabric. Intrinsic subtype annotation uses the following color scheme: red–Basal, blue–Luminal A, light blue–Luminal B, pink–Her2, green–normal-like, claudin-low–maroon, white–no annotation. Pink vertical highlight lines are included for aligning annotation with clusters.*

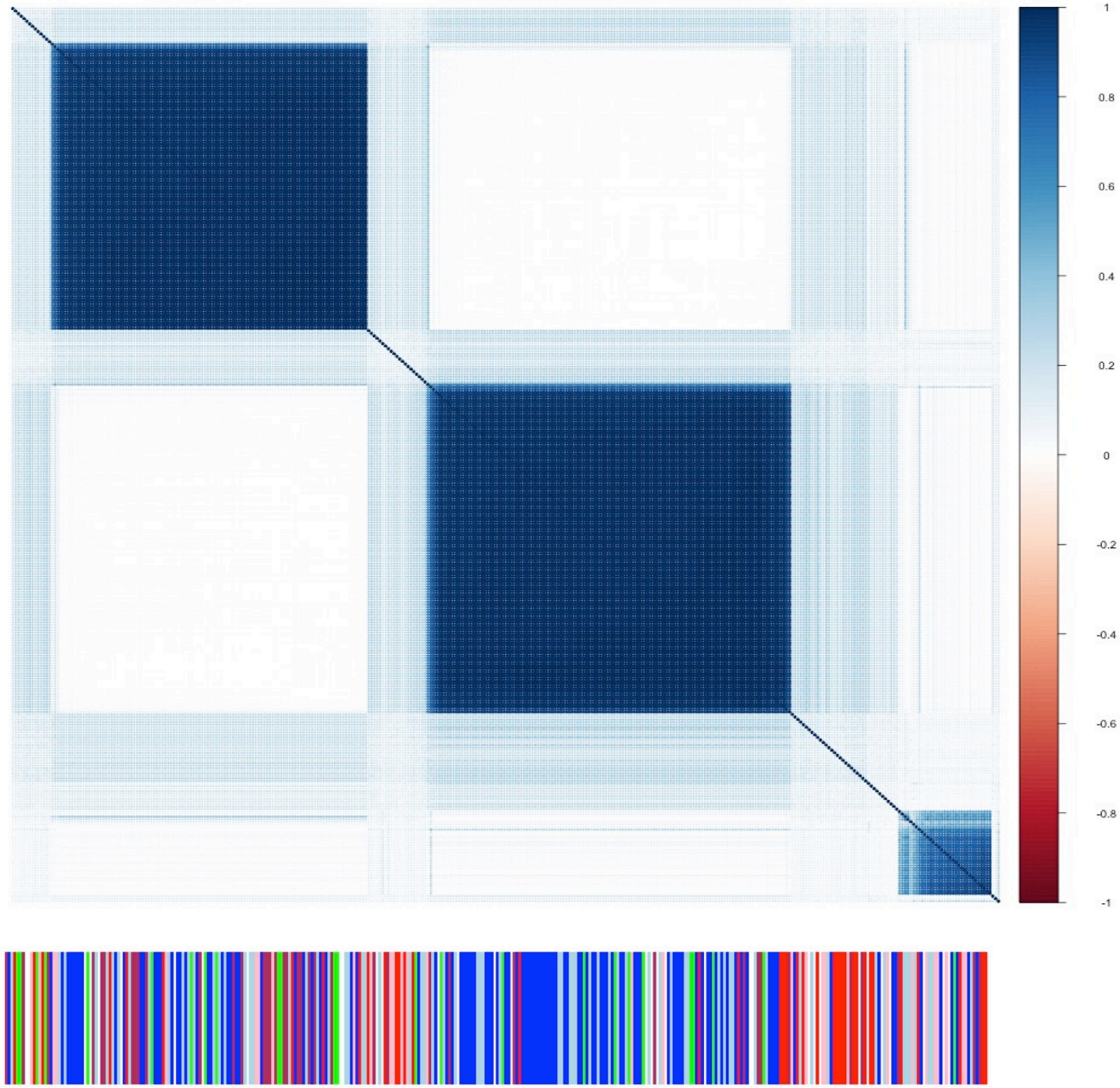


*Figure S11: CNA clustering posterior similarity matrix with subtype annotation of a sample of 350 subjects from Metabric. Intrinsic subtype annotation uses the following color scheme: red–Basal, blue–Luminal A, light blue–Luminal B, pink–Her2, green–normal-like, claudin-low–maroon, white–no annotation.*

### S6 TCGA Figures

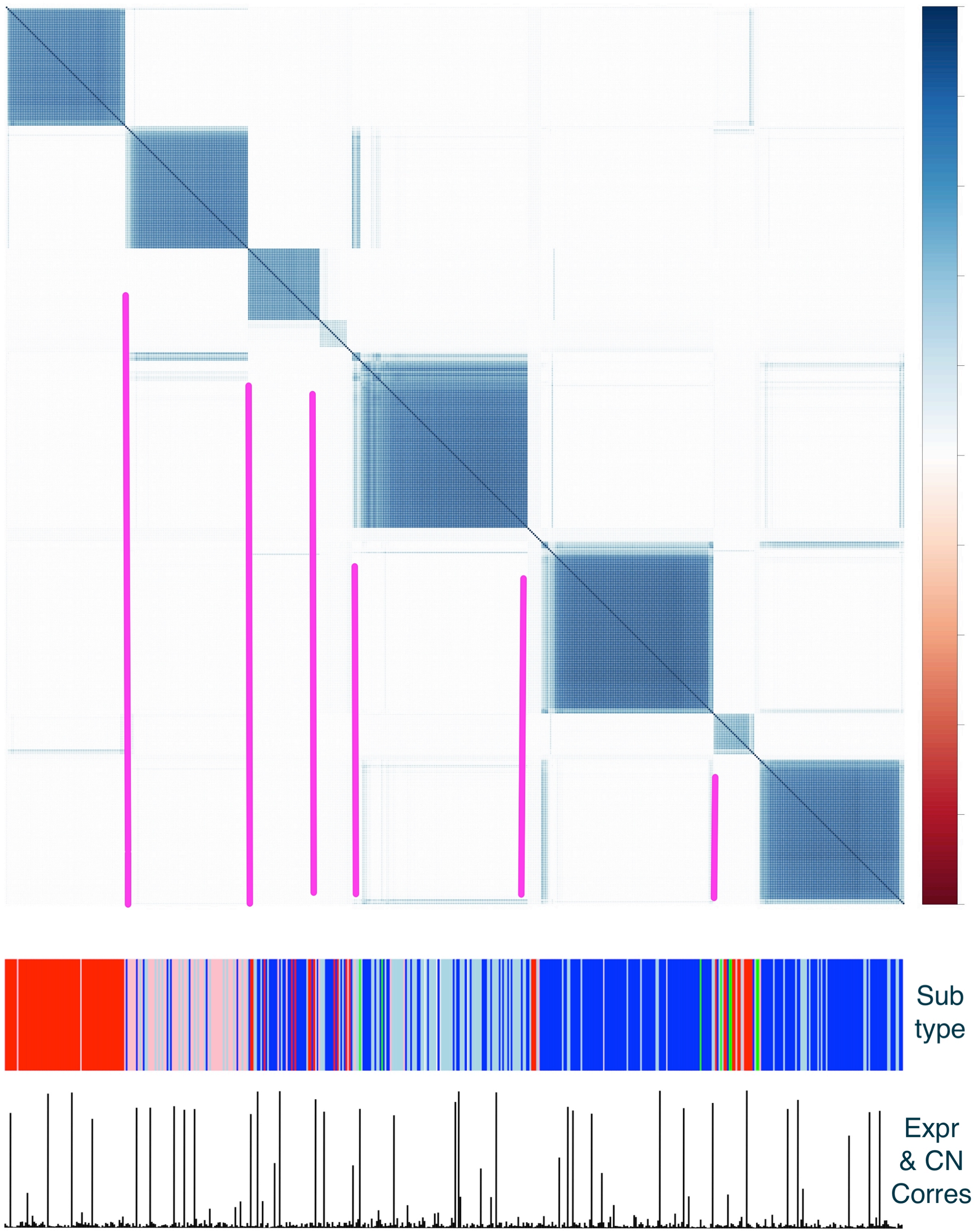


*Figure S12: TCGA expression clustering PSM with subtype annotation and expression-CNA cluster correspondence plotted below. Samples are ordered according to comprehensibility of expression clustering. Intrinsic subtype annotation uses the following color scheme: red–Basal, blue–Luminal A, light blue–Luminal B, pink–Her2, green–normal-like. Pink highlight lines included for aligning annotation with clusters.*

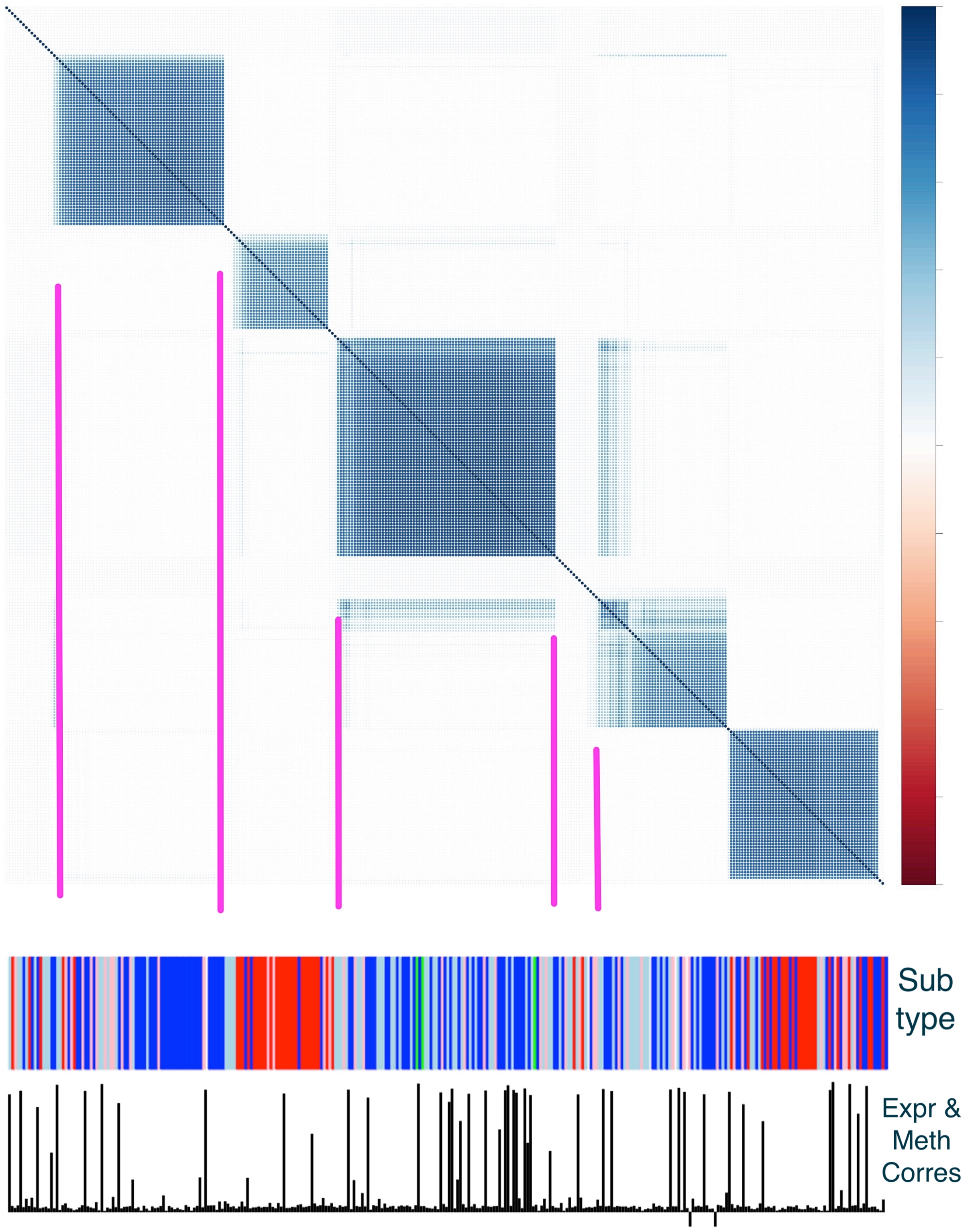


*Figure S14: TCGA methylation clustering PSM with subtype annotation and expression-methylation cluster correspondence plotted below. Samples are ordered according to comprehensibility of methylation clustering. Intrinsic subtype annotation uses the following color scheme: red–Basal, blue–Luminal A, light blue–Luminal B, pink–Her2, green–normal-like. Pink highlight lines included for aligning annotation with clusters.*

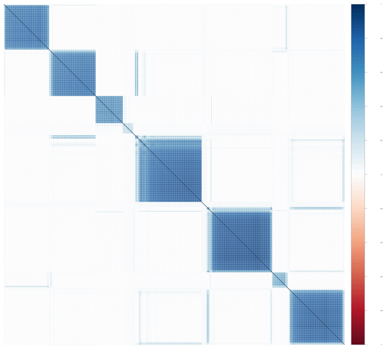

*(a) TCGA Expression PSM with corresponding ordering*

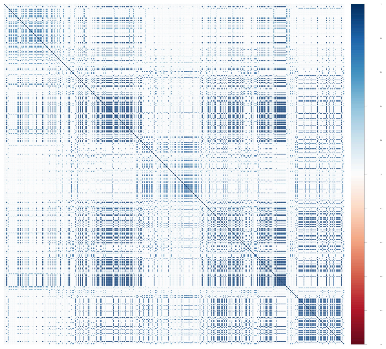

*(b) TCGA CNA PSM with expression ordering.*

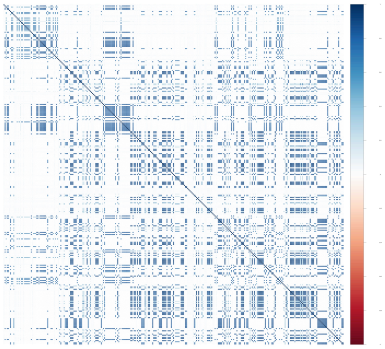

*(c) TCGA Methylation PSM with expression ordering.*

### S7 Supplementary Tables

*Table S1: Post-processed cluster labels for different feature draws from the expression dataset. We see an even higher degree of stability in methylation cluster labels across feature draws.*

| | clus_1 | clus_2 | clus_3 | clus_4 | clus_5 | clus_6 | clus_7 | clus_8 | clus_unknown | Sum |
|---|---|---|---|---|---|---|---|---|---|---|
| clus_1 | 0 | 16 | 13 | 0 | 0 | 0 | 0 | 0 | 0 | 29 |
| clus_2 | 2 | 0 | 1 | 0 | 0 | 1 | 0 | 34 | 0 | 38 |
| clus_3 | 0 | 0 | 0 | 54 | 0 | 0 | 0 | 3 | 0 | 57 |
| clus_4 | 0 | 0 | 0 | 6 | 0 | 0 | 19 | 2 | 0 | 27 |
| clus_5 | 6 | 0 | 0 | 3 | 0 | 0 | 1 | 0 | 3 | 13 |
| clus_6 | 13 | 0 | 0 | 0 | 0 | 0 | 0 | 0 | 0 | 13 |
| clus_7 | 9 | 0 | 0 | 1 | 0 | 3 | 7 | 2 | 0 | 22 |
| clus_8 | 0 | 0 | 0 | 0 | 8 | 45 | 0 | 0 | 1 | 54 |
| clus_unknown | 0 | 0 | 1 | 10 | 7 | 5 | 6 | 11 | 21 | 61 |
| Sum | 30 | 16 | 15 | 74 | 15 | 54 | 33 | 52 | 25 | 314 |

*Table S2: Post-processed cluster labels from parallel MCMC chains for the expression dataset. Since cluster labels are arbitrary, one could permute them so that the largest cells would fall along the diagonal. We see stability in cluster labels, indicating model convergence.*

| | clus_1 | clus_2 | clus_3 | clus_4 | clus_5 | clus_unknown | Sum |
|---|---|---|---|---|---|---|---|
| clus_1 | 0 | 0 | 37 | 1 | 11 | 1 | 50 |
| clus_2 | 29 | 1 | 1 | 1 | 9 | 1 | 42 |
| clus_3 | 2 | 57 | 1 | 0 | 2 | 5 | 67 |
| clus_4 | 0 | 0 | 0 | 0 | 0 | 28 | 28 |
| clus_5 | 0 | 4 | 0 | 109 | 28 | 17 | 158 |
| clus_unknown | 2 | 2 | 2 | 3 | 1 | 15 | 25 |
| Sum | 33 | 64 | 41 | 114 | 51 | 67 | 370 |

*Table S3: Post-processed cluster labels from parallel MCMC chains for the methylation dataset. Since cluster labels are arbitrary, one could permute them so that the largest cells would fall along the diagonal. We see a high degree of stability in cluster labels, indicating model convergence.*

| | clus_1 | clus_2 | clus_3 | clus_4 | clus_5 | clus_6 | clus_7 | clus_8 | clus_unknown | Sum |
|---|---|---|---|---|---|---|---|---|---|---|
| clus_1 | 0 | 0 | 0 | 0 | 0 | 0 | 32 | 1 | 5 | 38 |
| clus_2 | 26 | 0 | 0 | 2 | 0 | 0 | 0 | 0 | 1 | 29 |
| clus_3 | 1 | 0 | 0 | 72 | 0 | 0 | 0 | 9 | 0 | 82 |
| clus_4 | 2 | 0 | 0 | 0 | 0 | 1 | 0 | 34 | 0 | 37 |
| clus_5 | 0 | 0 | 0 | 0 | 2 | 49 | 0 | 0 | 1 | 52 |
| clus_6 | 0 | 0 | 15 | 0 | 0 | 2 | 0 | 0 | 0 | 17 |
| clus_7 | 0 | 15 | 0 | 0 | 0 | 0 | 0 | 0 | 0 | 15 |
| clus_unknown | 1 | 1 | 0 | 0 | 13 | 2 | 1 | 8 | 18 | 44 |
| Sum | 30 | 16 | 15 | 74 | 15 | 54 | 33 | 52 | 25 | 314 |

*Table S4: Post-processed cluster labels for the expression dataset tabulated against breast cancer subtypes. There is considerable association between the two sets of labels. Those observations with the more heterogeneous label ("clus_unknown") on the expression data source are more diffusely distributed across the subtypes than those with a clear cluster label.*

| | Basal | Her2 | LumA | LumB | Normal | Sum |
|---|---|---|---|---|---|---|
| clus_4 | 27 | 0 | 0 | 0 | 0 | 27 |
| clus_2 | 4 | 23 | 0 | 1 | 4 | 32 |
| clus_5 | 4 | 7 | 72 | 62 | 7 | 152 |
| clus_1 | 0 | 7 | 20 | 11 | 7 | 45 |
| clus_3 | 2 | 2 | 37 | 1 | 23 | 65 |
| clus_unknown | 11 | 8 | 9 | 13 | 8 | 49 |
| Sum | 48 | 47 | 138 | 88 | 49 | 370 |

*Table S5: Post-processed cluster labels for the methylation dataset tabulated against breast cancer subtypes. There is some association between the two sets of labels. There seems to be greater alignment between subtype and expression label than methylation label, which is expected since intrinsic subtype is defined with expression profiles.*

| | clus_1 | clus_2 | clus_3 | clus_4 | clus_5 | clus_6 | clus_7 | clus_unknown | Sum |
|---|---|---|---|---|---|---|---|---|---|
| LumB | 13 | 16 | 22 | 3 | 4 | 2 | 3 | 12 | 75 |
| Her2 | 12 | 3 | 4 | 1 | 12 | 4 | 2 | 5 | 43 |
| Normal | 2 | 1 | 3 | 5 | 12 | 7 | 1 | 3 | 34 |
| Basal | 1 | 1 | 1 | 1 | 16 | 4 | 0 | 17 | 41 |
| LumA | 10 | 8 | 52 | 27 | 8 | 0 | 9 | 7 | 121 |
| Sum | 38 | 29 | 82 | 37 | 52 | 17 | 15 | 44 | 314 |

*Table S6: We order observations according to sample-specific cluster correspondence and find enrichment in the invasive tumor state among those observations with greater expression-methylation cluster label sharing for different feature draws from the data (shown in (a) and (b)). The result suggests that there may be some degree of pan-genomic features acting in concert for tumors in a more advanced state.*

*a)*

| | DCIS | IDC |
|---|---|---|
| Most clust corresp samps | 0.024 | 0.976 |
| Least clust corresp samps | 0.048 | 0.952 |
| Marginal distribution | 0.154 | 0.846 |

*b)*

| | DCIS | IDC |
|---|---|---|
| Most clust corresp samps | 0.048 | 0.952 |
| Least clust corresp samps | 0.048 | 0.952 |
| Marginal distribution | 0.154 | 0.846 |

*Table S7: We order observations according to sample-specific cluster correspondence, and confirm using different feature draws enrichment in the HER2 intrinsic subtype among those observations with greater expression-CNA correspondence.*

*a)*

| | Basal | Her2 | LumA | LumB | Normal |
|---|---|---|---|---|---|
| Most clust corresp samps | 0.100 | 0.350 | 0.250 | 0.125 | 0.175 |
| Least clust corresp samps | 0.075 | 0.100 | 0.350 | 0.200 | 0.275 |
| Marginal distribution | 0.130 | 0.127 | 0.373 | 0.238 | 0.132 |

*b)*

| | Basal | Her2 | LumA | LumB | Normal |
|---|---|---|---|---|---|
| Most clust corresp samps | 0.100 | 0.425 | 0.075 | 0.250 | 0.150 |
| Least clust corresp samps | 0.075 | 0.125 | 0.350 | 0.200 | 0.250 |
| Marginal distribution | 0.130 | 0.127 | 0.373 | 0.238 | 0.132 |

*Table S8: We order observations according to sample-specific cluster correspondence, and find that there is slight enrichment in the HER2 subtype (a) and the LumA subtype (b) among those observations with greater cluster label sharing across the CNA-methylation and expression-methylation pairings, respectively. The result suggests that there is some degree of pan-genomic features acting in greater concert for tumors of the HER2 and LumA intrinsic subtypes.*

*a) CNA-methylation pairing*

| | Basal | Her2 | LumA | LumB | Normal |
|---|---|---|---|---|---|
| Most clust corresp samps | 0.000 | 0.267 | 0.400 | 0.267 | 0.067 |
| Least clust corresp samps | 0.333 | 0.133 | 0.267 | 0.200 | 0.067 |
| Marginal distribution | 0.130 | 0.127 | 0.373 | 0.238 | 0.132 |

*b) Expression-methylation pairing*

| | Basal | Her2 | LumA | LumB | Normal |
|---|---|---|---|---|---|
| Most clust corresp samps | 0.017 | 0.033 | 0.617 | 0.283 | 0.050 |
| Least clust corresp samps | 0.117 | 0.067 | 0.350 | 0.217 | 0.250 |
| Marginal distribution | 0.130 | 0.127 | 0.373 | 0.238 | 0.132 |

*Table S9: Ductal carcinoma is almost entirely found in the large CNA cluster across feature draws from the data (shown in (a) and (b)).*

*a)*

| | clus_1 | clus_2 | clus_3 | clus_4 | clus_unknown | Sum |
|---|---|---|---|---|---|---|
| DCIS | 3 | 42 | 0 | 0 | 3 | 48 |
| IDC | 62 | 160 | 2 | 2 | 64 | 290 |
| Sum | 65 | 202 | 2 | 2 | 67 | 338 |

*b)*

| | clus_1 | clus_2 | clus_3 | clus_unknown | Sum |
|---|---|---|---|---|---|
| DCIS | 3 | 1 | 40 | 4 | 48 |
| IDC | 63 | 24 | 129 | 74 | 290 |
| Sum | 66 | 25 | 169 | 78 | 338 |

*Table S10: We order observations according to sample-specific cluster correspondence and confirm using different feature draws enrichment in the DCIS tumor state among those observations with greater cluster label sharing across the expression-CNA pairing. The result suggests that there is some degree of expression-CNA features acting in greater concert for tumors that have not yet become invasive.*

*a)*

| | DCIS | IDC |
|---|---|---|
| Most clust corresp samps | 0.350 | 0.650 |
| Least clust corresp samps | 0.250 | 0.750 |
| Marginal distribution | 0.154 | 0.846 |

*b)*

| | DCIS | IDC |
|---|---|---|
| Most clust corresp samps | 0.425 | 0.575 |
| Least clust corresp samps | 0.275 | 0.725 |
| Marginal distribution | 0.154 | 0.846 |

*Table S11: There is a lack of the Basal intrinsic subtype among those observations with greater CNA-methylation sample-specific cluster correspondence across feature draws.*

*a)*

| | Basal | Her2 | LumA | LumB | Normal |
|---|---|---|---|---|---|
| Most clust corresp samps | 0.048 | 0.190 | 0.238 | 0.357 | 0.167 |
| Least clust corresp samps | 0.167 | 0.048 | 0.310 | 0.310 | 0.167 |
| Marginal distribution | 0.130 | 0.127 | 0.373 | 0.238 | 0.132 |

*b)*

| | Basal | Her2 | LumA | LumB | Normal |
|---|---|---|---|---|---|
| Most clust corresp samps | 0.024 | 0.214 | 0.286 | 0.381 | 0.095 |
| Least clust corresp samps | 0.167 | 0.048 | 0.310 | 0.310 | 0.167 |
| Marginal distribution | 0.130 | 0.127 | 0.373 | 0.238 | 0.132 |

*Table S12: Cross tabulation of intrinsic subtype and CNA clusters.*

| | clus_1 | clus_2 | clus_3 | clus_unknown | Sum |
|---|---|---|---|---|---|
| LumB | 17 | 15 | 22 | 27 | 81 |
| Her2 | 5 | 5 | 22 | 11 | 43 |
| Normal | 3 | 1 | 35 | 1 | 40 |
| Basal | 1 | 1 | 17 | 26 | 45 |
| LumA | 40 | 3 | 73 | 13 | 129 |
| Sum | 66 | 25 | 169 | 78 | 338 |

*Table S13: Cross tabulation of model defined expression subtype and tumor state. We observe significant enrichment in the DCIS tumor state in model defined cluster 1.*

| | clus_1 | clus_2 | clus_3 | clus_4 | clus_5 | clus_unknown | Sum |
|---|---|---|---|---|---|---|---|
| DCIS | 22 | 8 | 4 | 19 | 2 | 2 | 57 |
| IDC | 11 | 56 | 37 | 95 | 49 | 65 | 313 |
| Sum | 33 | 64 | 41 | 114 | 51 | 67 | 370 |

Christina Curtis, Sohrab P. Shah, Suet-Feung Chin, Gulisa Turashvili, Oscar M. Rueda, Mark J. Dunning, Doug Speed, Andy G. Lynch, Shamith Samarajiwa, Yinyin Yuan, Stefan Gräf, Gavin Ha, Gholamreza Haffari, Ali Bashashati, Roslin Russell, Steven McKinney, Carlos Caldas, Samuel Aparicio, Christina Curtis†, Sohrab P. Shah, Carlos Caldas, Samuel Aparicio, James D. Brenton, Ian Ellis, David Huntsman, Sarah Pinder, Arnie Purushotham, Leigh Murphy, Carlos Caldas, Samuel Aparicio, Carlos Caldas, Helen Bardwell, Suet-Feung Chin, Christina Curtis, Zhihao Ding, Stefan Gräf, Linda Jones, Bin Liu, Andy G. Lynch, Irene Papatheodorou, Stephen J. Sammut, Gordon Wishart, Samuel Aparicio, Steven Chia, Karen Gelmon, David Huntsman, Steven McKinney, Caroline Speers, Gulisa Turashvili, Peter Watson, Ian Ellis, Roger Blamey, Andrew Green, Douglas Macmillan, Emad Rakha, Arnie Purushotham, Cheryl Gillett, Anita Grigoriadis, Sarah Pinder, Emanuele di Rinaldis, Andy Tutt, Leigh Murphy, Michelle Parisien, Sandra Troup, Carlos Caldas, Suet-Feung Chin, Derek Chan, Claire Fielding, Ana-Teresa Maia, Sarah McGuire, Michelle Osborne, Sara M. Sayalero, Inmaculada Spiteri, James Hadfield, Samuel Aparicio, Gulisa Turashvili, Lynda Bell, Katie Chow, Nadia Gale, David Huntsman, Maria Kovalik, Ying Ng, Leah Prentice, Carlos Caldas, Simon Tavaré, Christina Curtis, Mark J. Dunning, Stefan Gräf, Andy G. Lynch, Oscar M. Rueda, Roslin Russell, Shamith Samarajiwa, Doug Speed, Florian Markowetz, Yinyin Yuan, James D. Brenton, Samuel Aparicio, Sohrab P. Shah, Ali Bashashati, Gavin Ha, Gholamreza Haffari, Steven McKinney, Anita Langerød, Andrew Green, Elena Provenzano, Gordon Wishart, Sarah Pinder, Peter Watson, Florian Markowetz, Leigh Murphy, Ian Ellis, Arnie Purushotham, Anne-Lise Børresen-Dale, James D. Brenton, Simon Tavaré, Carlos Caldas, and Samuel Aparicio. The genomic and transcriptomic architecture of 2,000 breast tumours reveals novel subgroups. *Nature*, April 2012. ISSN 0028-0836, 1476-4687. doi: 10.1038/nature10983. URL `http://www.nature.com/doifinder/10.1038/nature10983`.

David B. Dunson and Amy H. Herring. Bayesian latent variable models for mixed discrete outcomes. *Biostatistics*, 6(1):11–25, 2005.

Arno Fritsch and Katja Ickstadt. Improved criteria for clustering based on the posterior similarity matrix. *Bayesian Analysis*, 4(2):367–391, June 2009. ISSN 1936-0975. doi: 10.1214/09-BA414. URL `http://projecteuclid.org/euclid.ba/1340370282`.

Evelina Gabasova, John Reid, and Lorenz Wernisch. Clusternomics: Integrative context-dependent clustering for heterogeneous datasets. *PLOS Computational Biology*, 13(10): e1005781, October 2017. ISSN 1553-7358. doi: 10.1371/journal.pcbi.1005781. URL `http://dx.plos.org/10.1371/journal.pcbi.1005781`.

Kristoffer H. Hellton and Magne Thoresen. Integrative clustering of high-dimensional data with joint and individual clusters. *Biostatistics*, 17(3):537–548, July 2016. ISSN 1465-4644, 1468-4357. doi: 10.1093/biostatistics/kxw005. URL https://academic.oup.com/biostatistics/article-lookup/doi/10.1093/biostatistics/kxw005.

Christian Hennig. Breakdown points for maximum likelihood estimators of location-scale mixtures. *The Annals of Statistics*, 32(4):1313–1340, August 2004. ISSN 0090-5364. doi: 10.1214/009053604000000571. URL http://arxiv.org/abs/math/0410073. arXiv: math/0410073.

Katherine A. Hoadley, Christina Yau, Denise M. Wolf, Andrew D. Cherniack, David Tamborero, Sam Ng, Max D.M. Leiserson, Beifang Niu, Michael D. McLellan, Vladislav Uzunangelov, Jiashan Zhang, Cyriac Kandoth, Rehan Akbani, Hui Shen, Larsson Omberg, Andy Chu, Adam A. Margolin, Laura J. van't Veer, Nuria Lopez-Bigas, Peter W. Laird, Benjamin J. Raphael, Li Ding, A. Gordon Robertson, Lauren A. Byers, Gordon B. Mills, John N. Weinstein, Carter Van Waes, Zhong Chen, Eric A. Collisson, Christopher C. Benz, Charles M. Perou, and Joshua M. Stuart. Multiplatform Analysis of 12 Cancer Types Reveals Molecular Classification within and across Tissues of Origin. *Cell*, 158(4):929–944, August 2014. ISSN 00928674. doi: 10.1016/j.cell.2014.06.049. URL http://linkinghub.elsevier.com/retrieve/pii/S0092867414008769.

N. Huang, P. K. Shah, and C. Li. Lessons from a decade of integrating cancer copy number alterations with gene expression profiles. *Briefings in Bioinformatics*, 13(3): 305–316, May 2012. ISSN 1467-5463, 1477-4054. doi: 10.1093/bib/bbr056. URL https://academic.oup.com/bib/article-lookup/doi/10.1093/bib/bbr056.

Paul Kirk, Jim E. Griffin, Richard S. Savage, Zoubin Ghahramani, and David L. Wild. Bayesian correlated clustering to integrate multiple datasets. *Bioinformatics*, 28(24):3290–3297, December 2012. ISSN 1460-2059, 1367-4803. doi: 10.1093/bioinformatics/bts595. URL https://academic.oup.com/bioinformatics/article-lookup/doi/10.1093/bioinformatics/bts595.

Scott Kirkpatrick, C Daniel Gelatt, and Mario P Vecchi. Optimization by simulated annealing. *science*, 220(4598):671–680, 1983.

Daniel C. Koboldt, Robert S. Fulton, Michael D. McLellan, Heather Schmidt, Joelle Kalicki-Veizer, Joshua F. McMichael, Lucinda L. Fulton, David J. Dooling, Li Ding, Elaine R. Mardis, Richard K. Wilson, Adrian Ally, Miruna Balasundaram, Yaron S. N. Butterfield, Rebecca Carlsen, Candace Carter, Andy Chu, Eric Chuah, Noreen Dhalla, Carrie Hirst, Martin Hirst, Robert A. Holt, Darlene Lee, Haiyan I. Li, Michael Mayo, Richard A. Moore, Andrew J. Mungall, A. Gordon Robertson, Jacqueline E. Schein, Arash Shafiei, Payal Sipahimalani, Jared R. Slobodan, Dominik Stoll, Angela Tam, Nina Thiessen, Richard J. Varhol, Natasja Wye, Thomas Zeng, Yongjun Zhao, Inanc Birol, Steven J. M. Jones, Marco A. Marra, Andrew D. Cherniack, Gordon Saksena, Robert C. Onofrio, and Nam H. Pho. Comprehensive molecular portraits of human breast tumours. *Nature*, 490(7418): 61–70, September 2012. ISSN 0028-0836, 1476-4687. doi: 10.1038/nature11412. URL http://www.nature.com/doifinder/10.1038/nature11412.

Matthias Kormaksson, James G. Booth, Maria E. Figueroa, and Ari Melnick. Integrative Model-based clustering of microarray methylation and expression data. *The Annals of Applied Statistics*, 6(3):1327–1347, September 2012. ISSN 1932-6157. doi: 10.1214/11-AOAS533. URL http://projecteuclid.org/euclid.aoas/1346418585.

Vessela N. Kristensen, Ole Christian Lingjærde, Hege G. Russnes, Hans Kristian M. Vollan, Arnoldo Frigessi, and Anne-Lise Børresen-Dale. Principles and methods of integrative genomic analyses in cancer. *Nature Reviews Cancer*, 14(5):299–313, May 2014. ISSN 1474-175X, 1474-1768. doi: 10.1038/nrc3721. URL `http://www.nature.com/articles/nrc3721`.

R Lesurf, MR Aure, HH Mørk, V; Oslo Breast Cancer Research Consortium (OSBREAC) Vitelli, S Lundgren, AL Børresen-Dale, V Kristensen, F Wärnberg, M Hallett, and T Sørlie. Integrative clustering reveals a novel split in the luminal A subtype of breast cancer with impact on outcome. *Breast Cancer Research*, 19(1), December 2017. ISSN 1465-542X. doi: 10.1186/s13058-017-0812-y. URL `http://breast-cancer-research.biomedcentral.com/articles/10.1186/s13058-017-0812-y`.

Robert Lesurf, Miriam Ragle Aure, Hanne Håberg Mørk, Valeria Vitelli, Steinar Lundgren, Anne-Lise Børresen-Dale, Vessela Kristensen, Fredrik Wärnberg, Michael Hallett, Therese Sørlie, Torill Sauer, Jürgen Geisler, Solveig Hofvind, Elin Borgen, Anne-Lise Børresen-Dale, Olav Engebråten, Øystein Fodstad, Øystein Garred, Gry Aarum Geitvik, Rolf Kåresen, Bjørn Naume, Gunhild Mari Mælandsmo, Hege G. Russnes, Ellen Schlichting, Therese Sørlie, Ole Christian Lingjærde, Vessela Kristensen, Kristine Kleivi Sahlberg, Helle Kristine Skjerven, and Britt Fritzman. Molecular Features of Subtype-Specific Progression from Ductal Carcinoma In Situ to Invasive Breast Cancer. *Cell Reports*, 16 (4):1166–1179, July 2016. ISSN 22111247. doi: 10.1016/j.celrep.2016.06.051. URL `http://linkinghub.elsevier.com/retrieve/pii/S2211124716308051`.

Eric F. Lock and David B. Dunson. Bayesian consensus clustering. *Bioinformatics*, 29(20):2610–2616, October 2013. ISSN 1460-2059, 1367-4803. doi: 10.1093/bioinformatics/btt425. URL `https://academic.oup.com/bioinformatics/article-lookup/doi/10.1093/bioinformatics/btt425`.

Eric F. Lock, Katherine A. Hoadley, J. S. Marron, and Andrew B. Nobel. Joint and individual variation explained (JIVE) for integrated analysis of multiple data types. *The Annals of Applied Statistics*, 7(1):523–542, March 2013. ISSN 1932-6157. doi: 10.1214/12-AOAS597. URL `http://projecteuclid.org/euclid.aoas/1365527209`.

Qianxing Mo, Sijian Wang, Venkatraman E Seshan, Adam B Olshen, Nikolaus Schultz, Chris Sander, R Scott Powers, Marc Ladanyi, and Ronglai Shen. Pattern discovery and cancer gene identification in integrated cancer genomic data. *Proceedings of the National Academy of Sciences*, 110(11):4245–4250, 2013.

Qianxing Mo, Ronglai Shen, Cui Guo, Marina Vannucci, Keith S Chan, and Susan G Hilsenbeck. A fully Bayesian latent variable model for integrative clustering analysis of multi-type omics data. *Biostatistics*, 19(1):71–86, 2017.

Matahi Moarii, Valentina Boeva, Jean-Philippe Vert, and Fabien Reyal. Changes in correlation between promoter methylation and gene expression in cancer. *BMC Genomics*, 16(1), December 2015. ISSN 1471-2164. doi: 10.1186/s12864-015-1994-2. URL `http://www.biomedcentral.com/1471-2164/16/873`.

Aslaug Aamodt Muggerud, Michael Hallett, Hilde Johnsen, Kristine Kleivi, Wenjing Zhou, Simin Tahmasebpoor, Rose-Marie Amini, Johan Botling, Anne-Lise Børresen-Dale, Therese Sørlie, and Fredrik Wärnberg. Molecular diversity in ductal carcinoma *in situ* (DCIS) and early invasive breast cancer. *Molecular Oncology*, 4(4):

357–368, August 2010. ISSN 15747891. doi: 10.1016/j.molonc.2010.06.007. URL http://doi.wiley.com/10.1016/j.molonc.2010.06.007.

Simen Myhre, Ole-Christian Lingjaerde, Bryan T. Hennessy, Miriam R. Aure, Mark S. Carey, Jan Alsner, Trine Tramm, Jens Overgaard, Gordon B. Mills, Anne-Lise Børresen-Dale, and Therese Sørlie. Influence of DNA copy number and mRNA levels on the expression of breast cancer related proteins. *Molecular Oncology*, 7(3): 704–718, June 2013. ISSN 15747891. doi: 10.1016/j.molonc.2013.02.018. URL http://doi.wiley.com/10.1016/j.molonc.2013.02.018.

Dvir Netanely, Ayelet Avraham, Adit Ben-Baruch, Ella Evron, and Ron Shamir. Expression and methylation patterns partition luminal-A breast tumors into distinct prognostic subgroups. *Breast Cancer Research*, 18(1), December 2016. ISSN 1465-542X. doi: 10.1186/s13058-016-0724-2. URL http://breast-cancer-research.biomedcentral.com/articles/10.1186/s13058-016-0724-2.

Gro Nilsen, Knut Liestøl, Peter Van Loo, Hans Kristian Moen Vollan, Marianne B. Eide, Oscar M. Rueda, Suet-Feung Chin, Roslin Russell, Lars O. Baumbusch, and Carlos Caldas. Copynumber: Efficient algorithms for single-and multi-track copy number segmentation. *BMC genomics*, 13(1):591, 2012.

So Yeon Park, Mithat Gönen, Hee Jung Kim, Franziska Michor, and Kornelia Polyak. Cellular and genetic diversity in the progression of in situ human breast carcinomas to an invasive phenotype. *Journal of Clinical Investigation*, 120(2):636–644, February 2010. ISSN 0021-9738. doi: 10.1172/JCI40724. URL http://www.jci.org/articles/view/40724.

Joel S. Parker, Michael Mullins, Maggie C.U. Cheang, Samuel Leung, David Voduc, Tammi Vickery, Sherri Davies, Christiane Fauron, Xiaping He, Zhiyuan Hu, John F. Quackenbush, Inge J. Stijleman, Juan Palazzo, J.S. Marron, Andrew B. Nobel, Elaine Mardis, Torsten O. Nielsen, Matthew J. Ellis, Charles M. Perou, and Philip S. Bernard. Supervised Risk Predictor of Breast Cancer Based on Intrinsic Subtypes. *Journal of Clinical Oncology*, 27(8): 1160–1167, March 2009. ISSN 0732-183X, 1527-7755. doi: 10.1200/JCO.2008.18.1370. URL http://ascopubs.org/doi/10.1200/JCO.2008.18.1370.

R Core Team. *R: A Language and Environment for Statistical Computing*. R Foundation for Statistical Computing, Vienna, Austria, 2017. URL https://www.R-project.org/.

David J Reiss, Nitin S Baliga, and Richard Bonneau. Integrated biclustering of heterogeneous genome-wide datasets for the inference of global regulatory networks. *BMC Bioinformatics*, page 22, 2006.

Carlos E. Rodríguez and Stephen G. Walker. Label Switching in Bayesian Mixture Models: Deterministic Relabeling Strategies. *Journal of Computational and Graphical Statistics*, 23 (1):25–45, January 2014. ISSN 1061-8600, 1537-2715. doi: 10.1080/10618600.2012.735624. URL http://www.tandfonline.com/doi/abs/10.1080/10618600.2012.735624.

Ronglai Shen, Adam B. Olshen, and Marc Ladanyi. Integrative clustering of multiple genomic data types using a joint latent variable model with application to breast and lung cancer subtype analysis. *Bioinformatics*, 25(22):2906–2912, November 2009. ISSN 1460-2059, 1367-4803. doi: 10.1093/bioinformatics/btp543. URL https://academic.oup.com/bioinformatics/article-lookup/doi/10.1093/bioinformatics/btp543.

Ronglai Shen, Sijian Wang, and Qianxing Mo. Sparse integrative clustering of multiple omics data sets. *The Annals of Applied Statistics*, 7(1):269–294, March 2013. ISSN 1932-6157. doi: 10.1214/12-AOAS578. URL http://projecteuclid.org/euclid.aoas/1365527199.

Therese Sørlie, Charles M. Perou, Robert Tibshirani, Turid Aas, Stephanie Geisler, Hilde Johnsen, Trevor Hastie, Michael B. Eisen, Matt Van De Rijn, and Stefanie S. Jeffrey. Gene expression patterns of breast carcinomas distinguish tumor subclasses with clinical implications. *Proceedings of the National Academy of Sciences*, 98(19):10869–10874, 2001.

Therese Sørlie, Robert Tibshirani, Joel Parker, Trevor Hastie, James Stephen Marron, Andrew Nobel, Shibing Deng, Hilde Johnsen, Robert Pesich, and Stephanie Geisler. Repeated observation of breast tumor subtypes in independent gene expression data sets. *Proceedings of the National Academy of Sciences*, 100(14):8418–8423, 2003.

Nora K. Speicher and Nico Pfeifer. Integrating different data types by regularized unsupervised multiple kernel learning with application to cancer subtype discovery. *Bioinformatics*, 31(12):i268–i275, June 2015. ISSN 1367-4803, 1460-2059. doi: 10.1093/bioinformatics/btv244. URL https://academic.oup.com/bioinformatics/article-lookup/doi/10.1093/bioinformatics/btv244.

Wei Sun, Paul Bunn, Chong Jin, Paul Little, Vasyl Zhabotynsky, Charles M Perou, David Neil Hayes, Mengjie Chen, and Dan-Yu Lin. The association between copy number aberration, DNA methylation and gene expression in tumor samples. *Nucleic Acids Research*, February 2018. ISSN 0305-1048, 1362-4962. doi: 10.1093/nar/gky131. URL https://academic.oup.com/nar/advance-article/doi/10.1093/nar/gky131/4909987.

Nizar Touleimat and Jörg Tost. Complete pipeline for Infinium ® Human Methylation 450k BeadChip data processing using subset quantile normalization for accurate DNA methylation estimation. *Epigenomics*, 4(3):325–341, June 2012. ISSN 1750-1911, 1750-192X. doi: 10.2217/epi.12.21. URL http://www.futuremedicine.com/doi/10.2217/epi.12.21.

Yong Wang, Jill Waters, Marco L. Leung, Anna Unruh, Whijae Roh, Xiuqing Shi, Ken Chen, Paul Scheet, Selina Vattathil, Han Liang, Asha Multani, Hong Zhang, Rui Zhao, Franziska Michor, Funda Meric-Bernstam, and Nicholas E. Navin. Clonal evolution in breast cancer revealed by single nucleus genome sequencing. *Nature*, 512(7513):155–160, August 2014. ISSN 0028-0836, 1476-4687. doi: 10.1038/nature13600. URL http://www.nature.com/articles/nature13600.

Joe H. Ward. Hierarchical Grouping to Optimize an Objective Function. *Journal of the American Statistical Association*, 58(301):236, March 1963. ISSN 01621459. doi: 10.2307/2282967. URL http://www.jstor.org/stable/2282967?origin=crossref.

C S Wilhelm-Benartzi, D C Koestler, M R Karagas, J M Flanagan, B C Christensen, K T Kelsey, C J Marsit, E A Houseman, and R Brown. Review of processing and analysis methods for DNA methylation array data. *British Journal of Cancer*, 109(6):1394–1402, September 2013. ISSN 0007-0920, 1532-1827. doi: 10.1038/bjc.2013.496. URL http://www.nature.com/articles/bjc2013496.

T. A. Yap, M. Gerlinger, P. A. Futreal, L. Pusztai, and C. Swanton. Intratumor Heterogeneity: Seeing the Wood for the Trees. *Science Translational Medicine*, 4(127):127ps10–127ps10, March 2012. ISSN 1946-6234, 1946-6242. doi: 10.1126/scitranslmed.3003854. URL http://stm.sciencemag.org/cgi/doi/10.1126/scitranslmed.3003854.

Shihua Zhang, Chun-Chi Liu, Wenyuan Li, Hui Shen, Peter W. Laird, and Xianghong Jasmine Zhou. Discovery of multi-dimensional modules by integrative analysis of cancer genomic data. *Nucleic Acids Research*, 40(19):9379–9391, October 2012. ISSN 1362-4962, 0305-1048. doi: 10.1093/nar/gks725. URL https://academic.oup.com/nar/article-lookup/doi/10.1093/nar/gks725.
